# Meter-scale 2D clinostats uncover environmentally derived variation in tomato responses to simulated microgravity

**DOI:** 10.1101/2025.05.16.654566

**Authors:** Ashley N. Hostetler, Emily Kennebeck, Jonathan W. Reneau, Eva Birtell, Denise L. Caldwell, Anjali S. Iyer-Pascuzzi, Erin E. Sparks

## Abstract

Plants grown in spaceflight exhibit differences in physiology and morphology compared to those grown on Earth. While changes in gravity are a major environmental change, other space-related stressors make it difficult to identify the microgravity-specific responses. These knowledge gaps can be filled by ground-based microgravity simulators that randomize the perceived gravity vector, but these approaches have primarily been limited to smaller plants and seedlings. This study reports a set of meter-scale 2D clinostats that support the growth of plants beyond the seedling stage. Tomato plants were grown in five sequential trials under upright rotating control and clinorotated “simulated microgravity” conditions. We found that simulated microgravity impacted plant growth in each trial, but the response varied by trial. Analysis of environmental co-variates across trials revealed that temperature significantly contributed to variation in plant growth. Further, our results show that moderate heat stress can promote plant growth under simulated microgravity. Thus, this work demonstrates the potential of meter-scale clinostats to uncover interactions between the environmental and simulated microgravity, which alter plant growth.

## 1. Introduction

Sustaining life beyond Earth requires a deep understanding of how living organisms grow and adapt to lower gravity environments. For plants, spaceflight studies have shown a multitude of impacts on growth and development, which vary between studies (Nie et al., 2025; Wheeler, 2017). Differences in hardware, radiation, and CO_2_, introduce a significant source of variation for spaceflight experiments (De Micco et al., 2023). This variation makes it difficult to parse out plant responses to individual spaceflight components (Burgner et al., 2019, 2020). To overcome this limitation, spaceflight experiments have been complemented by ground-based approaches that use microgravity simulators to understand the impact of microgravity in different environmental contexts (Ferranti et al., 2020).

Pervasive among space environment simulators are microgravity simulators, which include 1D, 2D and 3D clinostat systems and random position machines (Hasenstein and Loon, 2015). These approaches randomize the gravity vector and prevent statoliths from sedimenting. Statoliths are the primary gravity sensing mechanism of plants and sediment in the direction of gravity. When they are prevented from sedimenting, this simulates a microgravity environment. These systems are predominantly designed for smaller plants grown on plates (Hoson et al., 1992; Ngwoke et al., 2023; Totsline et al., 2024). Although these experiments provide vital insights on plant germination and initial growth, there are only a few reports on the effects of microgravity beyond the seedling stage. One study using the model plant *Arabidopsis thaliana* showed that simulated microgravity inhibited plant reproduction to varying degrees among different ecotypes but had no impact on the earlier developmental stages (Miyamoto et al., 1999). A separate study grew the dwarf tomato cultivar, Micro-Tom, to maturity on a clinostat and showed inhibited vegetative (i.e., leaf area; dry weight) and reproductive (i.e., fruit yield; fruit weight) characteristics (Colla et al., 2007). These studies emphasize the importance of studying the impact of simulated microgravity beyond the seedling stage. However, there are limited 2D clinostat systems that allow crop plant growth beyond the seedling stage to enable these studies.

In this study, we designed and built two meter-scale 2D clinostats (1.5748 m × 2.1336 m × 1.5748 m), one rotating perpendicular to gravity (termed simulated microgravity) and one rotating parallel to gravity (control). We used these systems to test the hypothesis that simulated microgravity (i.e., randomizing the gravity vector) impacts the vegetative growth and development of tomato plants beyond the seedling stage. Across five trials, changes in plant growth under simulated microgravity were observed in each trial, however the direction of change (whether inhibited or enhanced) was impacted by the environmental conditions of each trial period. Our results indicate that moderate heat stress can alleviate the negative impacts of simulated microgravity on tomato growth. Collectively, these results demonstrate utility of meter-scale clinostats to uncover the role of environmental cues in the plant response to simulated microgravity.

## 2. Materials and Methods

### 2.1. Clinostat Design and Construction

Two clinostats were designed and constructed: one for upright rotational control and one for simulated microgravity. A full parts list can be found in **Table S1**. For both clinostats, frames were constructed from 80-20 T-slot aluminum extrusion framing. Attachments between extrusions were achieved by L-brackets. In total, each frame measured 1.5748 m (62 in) x 2.1336 m (84 in) x 1.5748 m (62 in). A center shaft of 50.80 mm (2 in) diameter aluminum round stock with a wall thickness of 9.525 mm (0.375 in) was run through the center of the frames. The top of the frame contained light-emitting diode (LED) light bars, each attached to a heatsink. Four rhizoboxes were attached to the center shaft at 16 in (40.64 cm), and each rhizobox was divided in half. Rotation was achieved by pairing a 12-volt DC motor to a spur gear box designed to have a sub 5 revolutions per minute (RPM) spin rate. Rotation speed was set to approximately 3 RPM to minimize shear stress (Lyon, 1970).

### 2.2. Experimental Design

Five trials were conducted between 29 January 2024 and 12 June 2024 at the University of Delaware, Newark, Delaware, USA. Each trial had a duration of 29 days from seed sterilization to harvest. Trials were named for their month of harvest (February, March, April, May, and June). Within each trial, two tomato (*Solanum lycopersicum*) cultivars, Moneymaker and Hawaii7996, were grown with four plants each under upright control and simulated microgravity conditions. Each cultivar and gravity combination was further split into *Fusarium oxysporum* or mock treatments. Treatments were randomly assigned to rhizoboxes within each trial, and each rhizobox contained the same *F. oxysporum* treatment (inoculation or mock) to limit any potential cross-contamination. No *F. oxysporum* infection or disease symptoms were observed in either upright control or simulated microgravity plants, and there was a complete overlap of traits (**Figure S1**). Therefore, *F. oxysporum* was disregarded as a factor in this study.

### 2.3. Seed Sterilization and Germination

Seeds were sterilized with a 50% household bleach solution (1:1 solution of 8.3% sodium hypochlorite and water) and washed in ultrapure water before overnight imbibition at 4 °C. Seeds were planted in a 72-cell tray filled with soil (LM-1 Germination Mix, Lambert) and saturated with water. To compensate for different germination times, Moneymaker seeds were started on Day 1 and Hawaii7996 seeds were started on Day 3. Trays were placed in a growth chamber (CU-22L, Percival Scientific) with fluorescent and white LED bulbs set to a 16 hr photoperiod and 198 µmol/m^2^/s. Day/night temperatures were set to 28 °C/25 °C. Seedlings were watered with a two-part nutrient solution (5-12-26 Part A and Cal Nit Part B, JR Peters Inc) on Day 9. On Day 13, the seedling trays were transferred to the greenhouse.

### 2.4. Plant Growth on Clinostats

The greenhouse had high-pressure sodium lamps providing supplemental lighting and day/night temperatures of 26.7 °C/21.1 °C. On Day 14, plants were transferred to rhizoboxes, that were divided into two panels. Each panel was filled with 900 g of soil. The top 1/3 of each panel was saturated with 250 mL of the same nutrient solution as seedlings and 300 mL autoclaved half-strength Potato Dextrose Broth or 300 mL half-strength Potato Dextrose Broth with 2.4 × 10^6^ – 10.5 × 10^6^ *F. oxysporum* spores per mL. No *F. oxysporum* infection or disease symptoms were observed, so *F. oxysporum* was disregarded as a factor in this study and all plants were analyzed together. The lower 2/3 of each panel was saturated with an additional 400 mL of nutrient solution. A piece of rockwool (2.54 cm × 3.81 cm × 3.175 cm, Gro-Slab) was cut in half, placed at the hypocotyl-root junction of a seedling, and inserted into the top of the panel. Rhizoboxes were closed and secured to the clinostat frames. Data was collected on Day 29, after 15 days of growth on the clinostats.

### 2.5. Plant Trait Evaluation

On Day 29, the following plant traits were measured: apical bud height (cm), shoot fresh mass (g), and average stem diameter (mm). Shoot dry mass (g) was measured after 7 days of drying in a 65 °C incubator. Apical bud height was measured from the rockwool to the tallest apical bud. Two stem diameter measurements were taken below the first stem node above the rockwool one diameter parallel relative to the width of the rhizobox and one perpendicular relative to the width of the rhizobox. These values were then averaged. Root images were obtained with a Canon EOS Rebel T6 and segmented with RootPainter (Smith et al., 2022). Data was removed for any plant where there was evidence of soil disruption (**Figure S2**). In RootPainter, the foreground and background of each image were manually annotated to train the model. After training, the model was applied to all images and root length data was extracted. For a small subset of images (n=10), although shoots were visible and appeared healthy, roots were not visually identifiable and segmentation failed. For these plants, a value of 0 was used to indicate no visible root system.

### 2.6. Environmental Data

A HOBO Temperature/Relative Humidity Datalogger (MX1101, Onset) recorded greenhouse conditions at 10 min intervals. The average temperature and average relative humidity were calculated across the duration of each trial. The average daytime and nighttime temperatures were calculated considering a 16 hr photoperiod. Hourly solar radiation data was obtained from the Delaware Environmental Observing System (DEOS; www.deos.udel.edu) using the Newark, DE-Ag Farm station. The cumulative solar radiation was calculated by summing the data across the duration of each trial.

### 2.7. Statistical Analyses

Statistical analyses were completed in R *version* 4.4.0 (Team, 2013) and figures were generated with the ggplot2 package *version* 3.5.1 (Wickham, 2009) and the cowplot package *version* 1.1.3 (Wilke, 2024). To reduce the dimensionality of the data, identify traits that best explain the variability in the data, and visualize clustering of data, principal component analyses (PCAs) were used. For PCAs, the data was centered and scaled, and the following phenotypes were included: apical bud height, average stem diameter, shoot fresh mass, shoot dry mass, and root length. Two different PCA were considered (1) a single PCA with traits from all trials combined, and (2) five individual PCAs (one for each trial). To visualize separation of groups, convex hulls based on meta-data were generated around samples.

To identify the impact of environment on plant traits, two linear mixed-effects models were fit for each trait using the lme4 package *version* 1.1.37 (Bates et al., 2014). The first model was the baseline model, which included cultivar and gravity condition as main effects and trial as a random effect. The second model was the environmental model, which included cultivar and gravity condition as main effects, trial as a random effect, and scaled environmental data as covariates. For each trait, the baseline model was compared to the environmental model using a likelihood ratio test, which determined if including covariates significantly improved the model.

To identify the impact of cultivar and gravity condition on plant traits, linear mixed-effects models were used for each trait within each trial. Each model included cultivar, gravity condition, and their interaction as main effects. If residuals were not normally distributed, a Tukey’s ladder of powers (rcompanion package *version* 2.5.0 (Mangiafico, 2021)) transformation was used. When a main effect was found to be significant (*P*<0.05), pairwise contrasts were computed using the emmeans package *version* 1.11.1 (Lenth, 2025).

To identify the impact of trial on plant traits, linear mixed-effects models were used for each trait within a gravity condition. Each model included cultivar and trial as main effects. If residuals were not normally distributed, a Tukey’s ladder of powers transformation was used. An analysis of variance (ANOVA) was performed. A post-hoc Tukey honest significant difference test was used to test all pairwise comparisons when a main effect was found to be significant (*P*<0.05).

## 3. Results

A set of meter-scale 2D clinostats were built to allow paired plant growth in simulated microgravity beyond the seedling stage (**Figure 1A-B**). One clinostat rotated parallel to the gravity vector (control; **Figure 1C**), while a second clinostat rotated perpendicular to the gravity vector (simulated microgravity; **Figure 1D**). These clinostats contained modifications to enable plant growth beyond the seedling stage. Each clinostat had an integrated LED panel positioned 95 cm from the clinostat surface, to limit phototropism competition. Rhizoboxes were used for plant growth (**Figure 1B**), which enabled the visualization of the root systems after the experiment.

**Figure 1.**
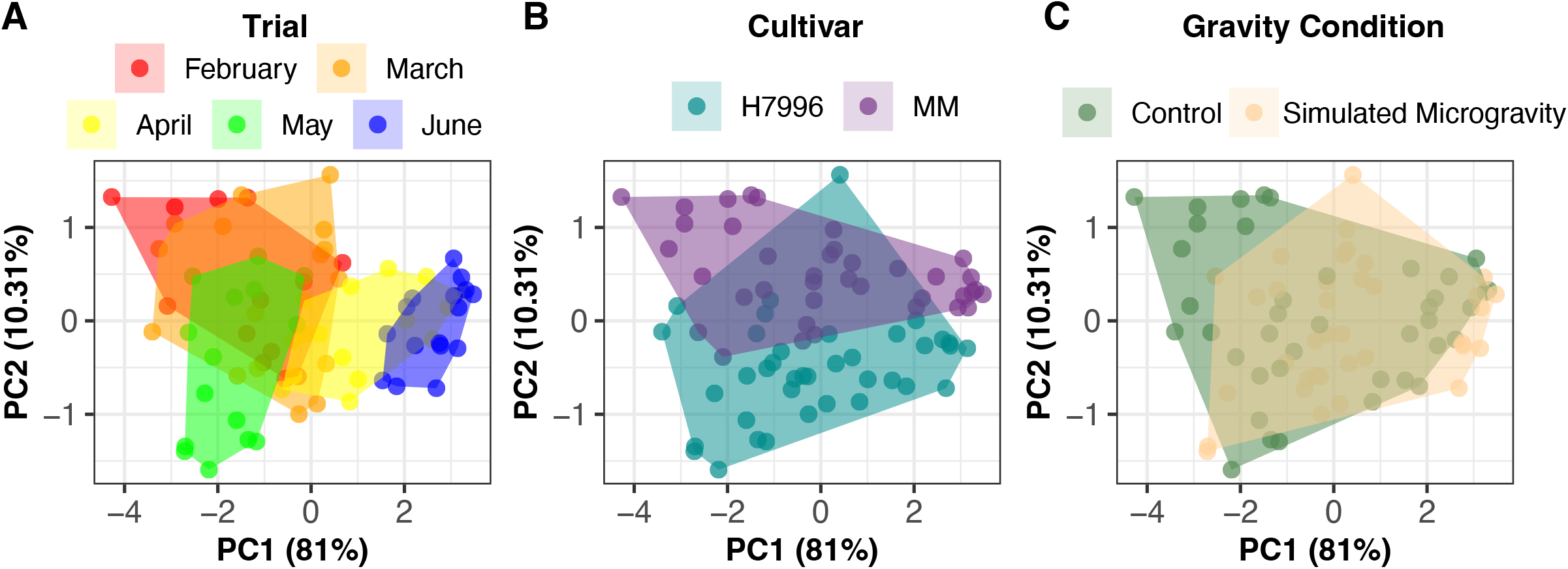
Meter-Scale 2D Clinostats. (A) Design of a modular frame for a 2D clinostat that supports plant growth beyond the seedling stage. (B) Soil-filled rhizoboxes were attached to the frame for rotation. (C) Clinostat in upright control position. (D) Clinostat in simulated microgravity position. (E) Tomato plant growing in clinostat.

Using the meter-scale clinostats, five consecutive trials of two tomato cultivars and two gravity conditions were conducted. A PCA analysis of plant traits (apical bud height, aboveground fresh mass, aboveground dry mass, shoot diameter, and total root length) after 15 days of rotation showed PC1 explained 81% of the variation and PC2 explained 10.31% of the variation **(Figure 2, Table S2)**. The variation across PC1 was best explained by trial (**Figure 2A**), whereas the variation across PC2 was best explained by cultivar (**Figure 2B**). There was minimal clustering by gravity condition (**Figure 2C**).

**Figure 2.**
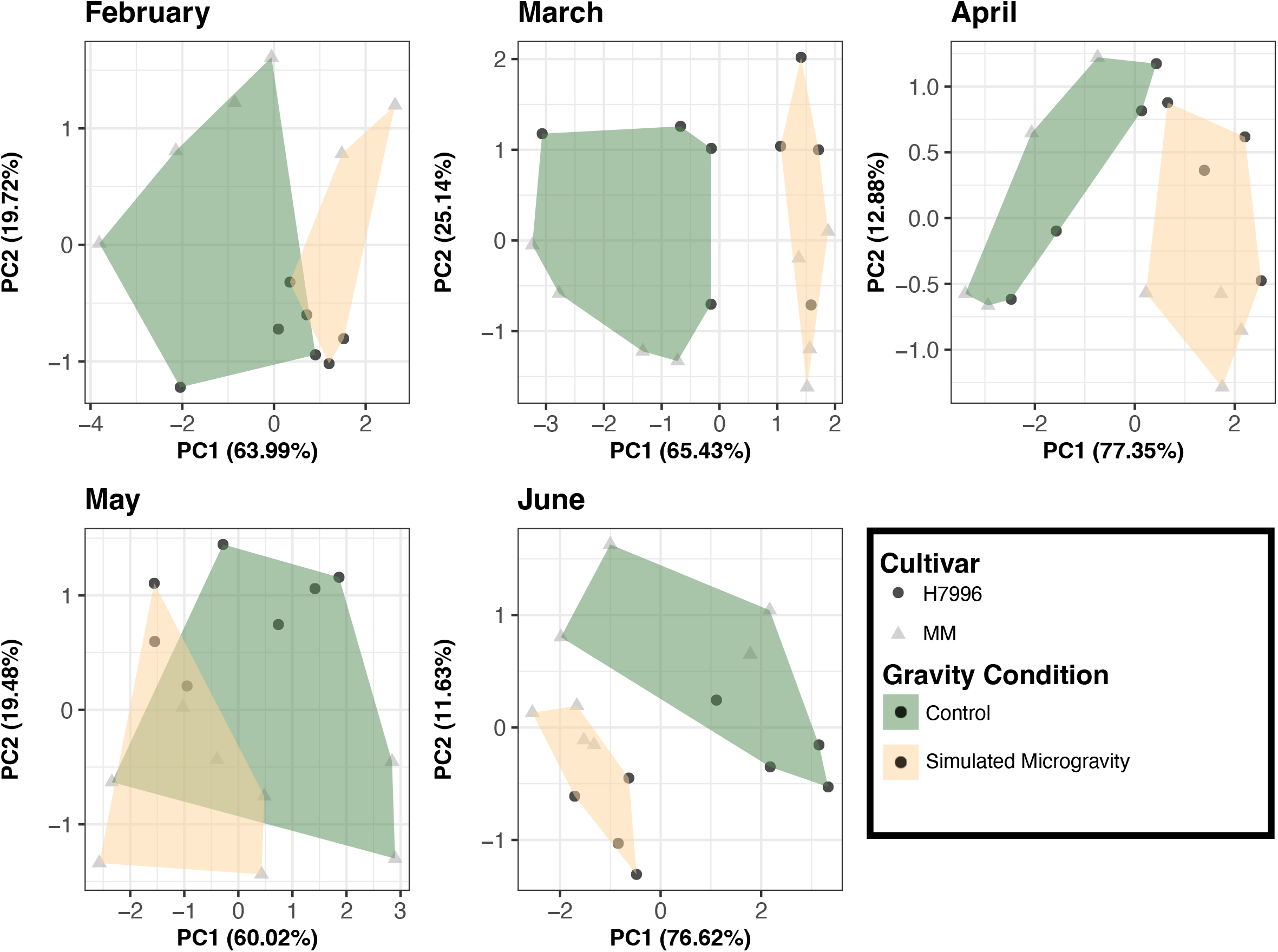
PCA of traits shows overlap by trial, cultivar, and gravity condition. PC1 and PC2 explain approximately 91% of the phenotypic variance. (A) Variation across PC1 is best explained by trial. (B) Variation across PC2 is best explained by cultivar. (C) There is minimal separation of data by gravity conditions. (A-C) Each point represents a single plant from five trials.

Given that trial best explained the variation across PC1, we separated the data by trial for additional analyses. When a PCA was applied to data within each trial, PC1 explained between 60–77% of the variation and PC2 explained 11–25% of the variation **(Figure 3, Table S3)**. Together, these two PCs explain between 79–90% of the variation in the plant traits. Variation across PC1 for all trials was best explained by gravity condition (**Figure 3**). Whereas variation across PC2 was best explained by cultivar (**Figure S3, Table S4**). These data demonstrate that, although the trial introduced variation in the data, within a trial, plant traits are separable by both gravity condition and cultivar.

**Figure 3.**
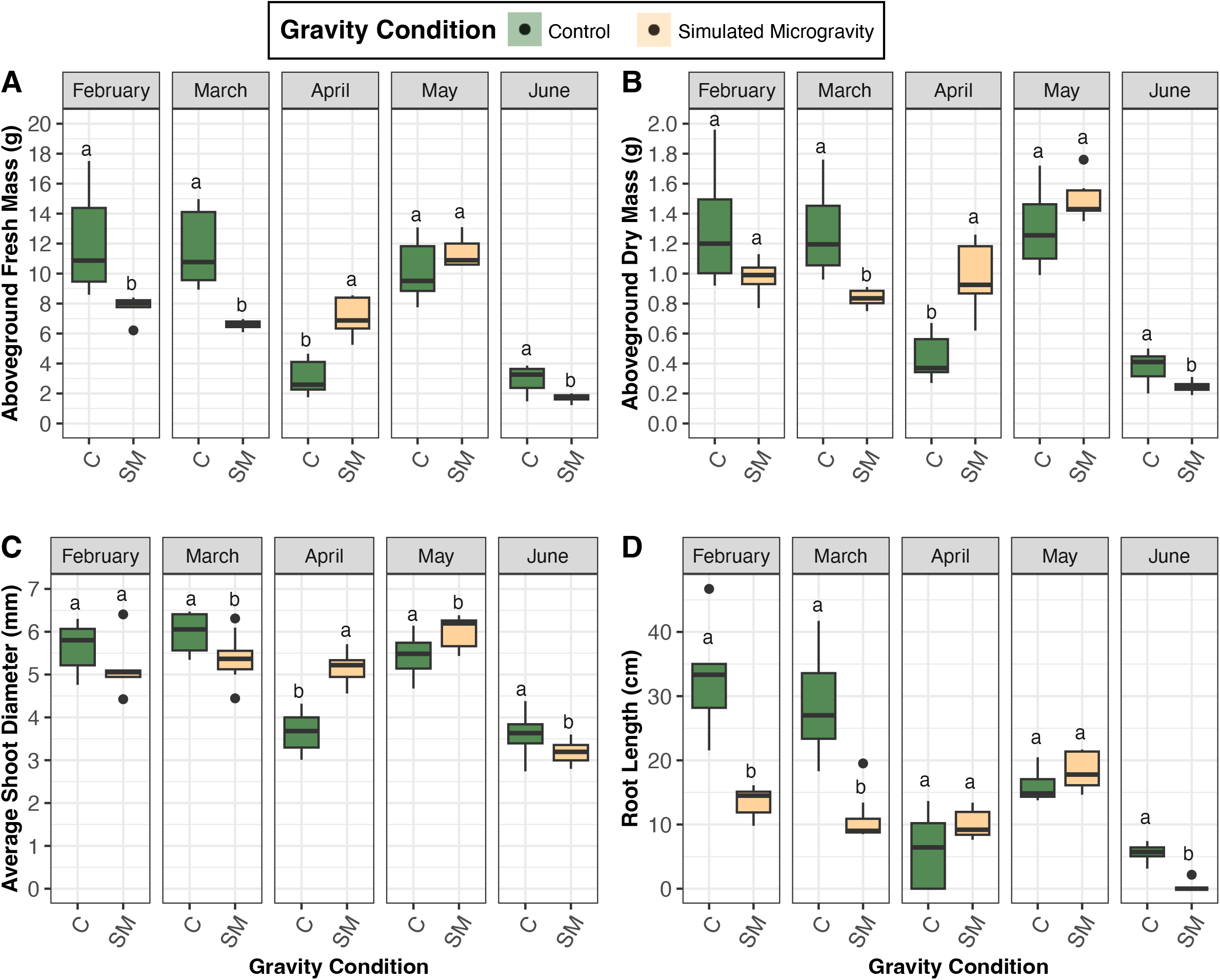
PCA of traits by trial reveals separation by cultivar and gravity condition. Within each trial, principal component analysis (PCA) of plant traits shows clear separation by gravity condition along PC1 and by cultivar along PC2. Across trial, PC1 accounts for 60-77% of the phenotypic variation, and PC2 explains 11-25%.

In considering the factors that could have led to trial effects, we hypothesized that the environment was a significant driver. The size of the clinostats necessitated the use of a greenhouse, and the long duration of each trial led to the five trials spanning almost six months. Analysis of environmental data showed that there was variation in the average temperature, average relative humidity, and cumulative solar radiation between trials (**Table S5**). To determine if inclusion of environmental data explained the trait variation among trials, we applied a likelihood ratio test. For all plant traits, inclusion of all three environmental covariates significantly improved the model (*P*<0.05) and explained additional variation (**Table 1; Table S6**). When assessing individual environmental covariates, average temperature and cumulative solar radiation improved the model, whereas average relative humidity had no effect (**Table 1; Table S6**). The average daily (**Figure S4A**) and hourly temperatures (**Figure S4B**) showed significant variation among the trials. Therefore, the average daytime and average nighttime temperatures were also tested as covariates in the model. The inclusion of either average daytime or average nighttime temperature did not improve the the model fit relative to the average daily temperature model (**Table S6**), suggesting that differences are primarily driven by overall temperature rather than distinct diurnal temperature effects. These results show that environmental changes across the six-month period contributed to differences among trials.

**Table 1.** Summary of likelihood ratio test evaluating model performance with the inclusion of environmental covariates.

|  | *P* (Base - Env) | *P* (Base - Temp) | *P* (Base - RH) | *P* (Base - Solar) |
| --- | --- | --- | --- | --- |
| **Apical Bud Height** | ** | n.s | n.s | ** |
| **Shoot Fresh Mass** | *** | ** | n.s | ** |
| **Shoot Dry Mass** | *** | * | n.s | ** |
| **Shoot Diameter** | *** | ** | n.s | *** |
| **Root Length** | *** | *** | n.s | *** |

\*\*\* $P < 0.001$
\*\* $P < 0.05$
\* $P < 0.1$

To quantify the impact of simulated microgravity on plant traits independent of environmental variation, data were evaluated within each trial to determine the effects of cultivar, gravity condition, and their interaction. Across all trials, cultivar impacted apical bud height (*P*<0.05), but not shoot fresh mass, shoot dry mass, shoot diameter, or root length (*P*>0.05) (**Table S7**). In all cases, Moneymaker had a shorter apical bud height than Hawaii7996 (**Figure S5, Table S8**). This is consistent with the slower growth rate in Moneymaker compared to Hawaii7996. In contrast, gravity condition significantly affected shoot fresh mass, shoot dry mass, shoot diameter, and root length (*P*<0.05), but not apical bud height (*P*>0.05) (**Table S7**). There were no consistent interaction effects between cultivar and gravity condition.

Analysis of the trial-specific data showed variable plant responses to simulated microgravity (**Figure 4, Table S8**). Specifically, in April and May trials, simulated microgravity positively affected all traits, while in February, March, and June, simulated microgravity negatively affected all traits (**Figure 4, Table S8**). When considering if plant water status was impacting these results, findings again show that the response of percent dry mass to simulated microgravity was trial-specific, with some trials showing an increase in percent dry mass and others showing a decrease in percent dry mass (**Tables S7-S8, Figure S6**). However, we noted substantial variation in the scale of data across trials, which was evident under both control and simulated microgravity conditions.

**Figure 4.**
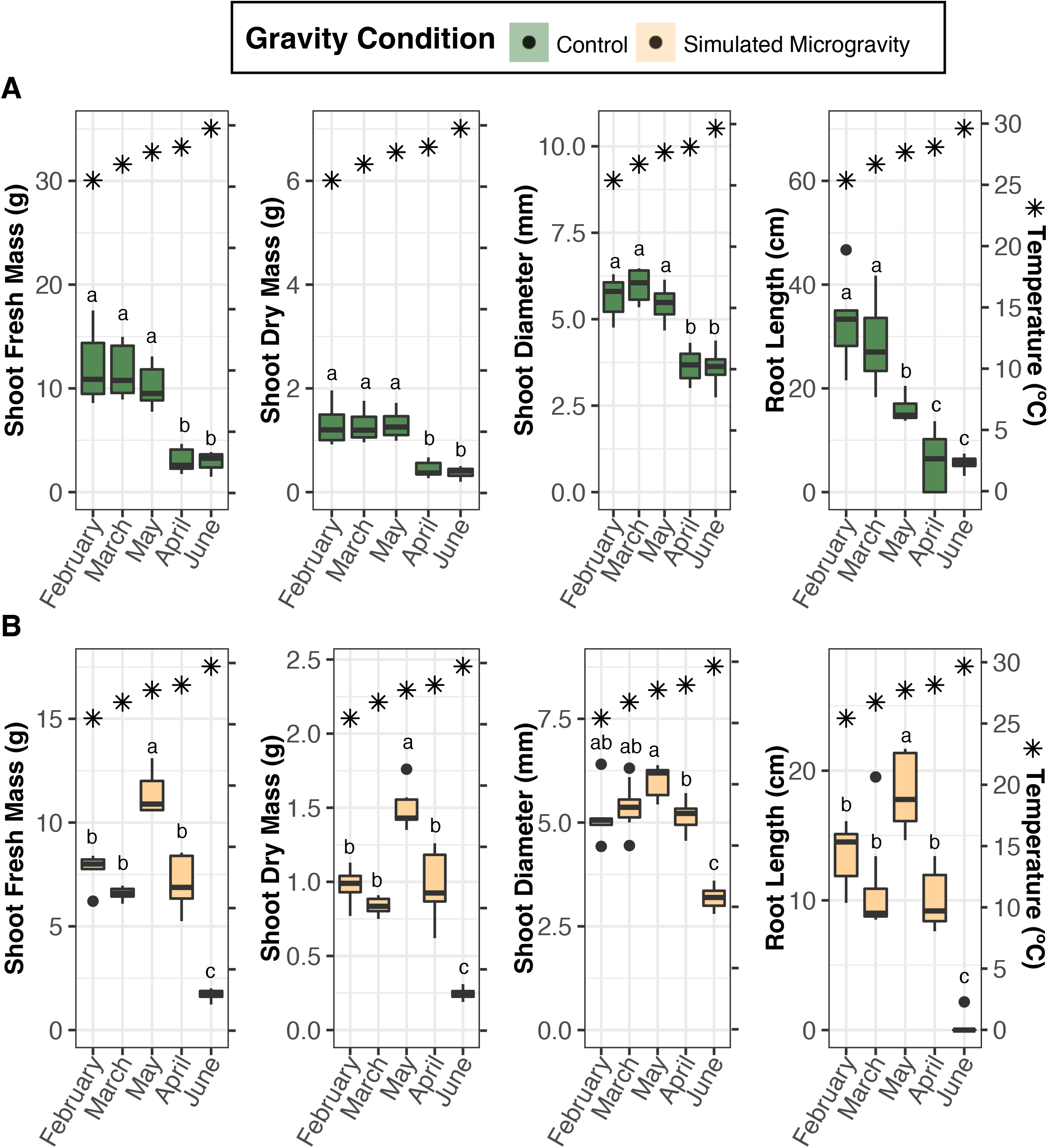
Plant responses to simulated microgravity were variable across trials. For (A) aboveground fresh mass, (B) aboveground dry mass, (C) shoot diameter, and (D) root length simulated microgravity significantly impacted plant traits (*P*<0.05), however the directional impact was variable depending on trial. For all traits (A-D), simulated microgravity negatively impacted traits in February, March, and June, while it positively impacted traits in April and May. Gravity conditions that share a letter are not significantly different within a trial (*P*>0.05). Black dots indicate outliers. C = control; SM = simulated microgravity

To assess the differences among upright control conditions, we identified three distinct response groups (**Figure 5A, Table S9-10**). These groupings showed reduced plant growth corresponding with increased temperature (**Figure 5A**) and cumulative solar radiation (**Figure S7**). Since increased solar radiation should increase plant growth, we hypothesized that the differences in control responses were due to temperature differences between trials. When we calculated average daytime temperatures they were in the range of standard tomato greenhouse production (February 27.90 °C +/- 4.11 and March 29.44 °C +/- 4.95), and the control plants had the highest shoot and root growth among control plants from all trials (**Figure 5A**). As the temperature increased (May 30.74 °C +/- 3.56), the control plants had comparable shoot growth to standard temperature conditions but reduced root growth (**Figure 5A**). When the temperatures continue to increase (April 31.32 °C +/- 3.95 and June 33.13 °C +/- 3.69), there was a reduction in both shoot and root growth in control plants compared to control plants from other trials (**Figure 5A**). These response groupings are notable in that they do not follow sequential trial order, as May had lower temperatures than April.

**Figure 5.**
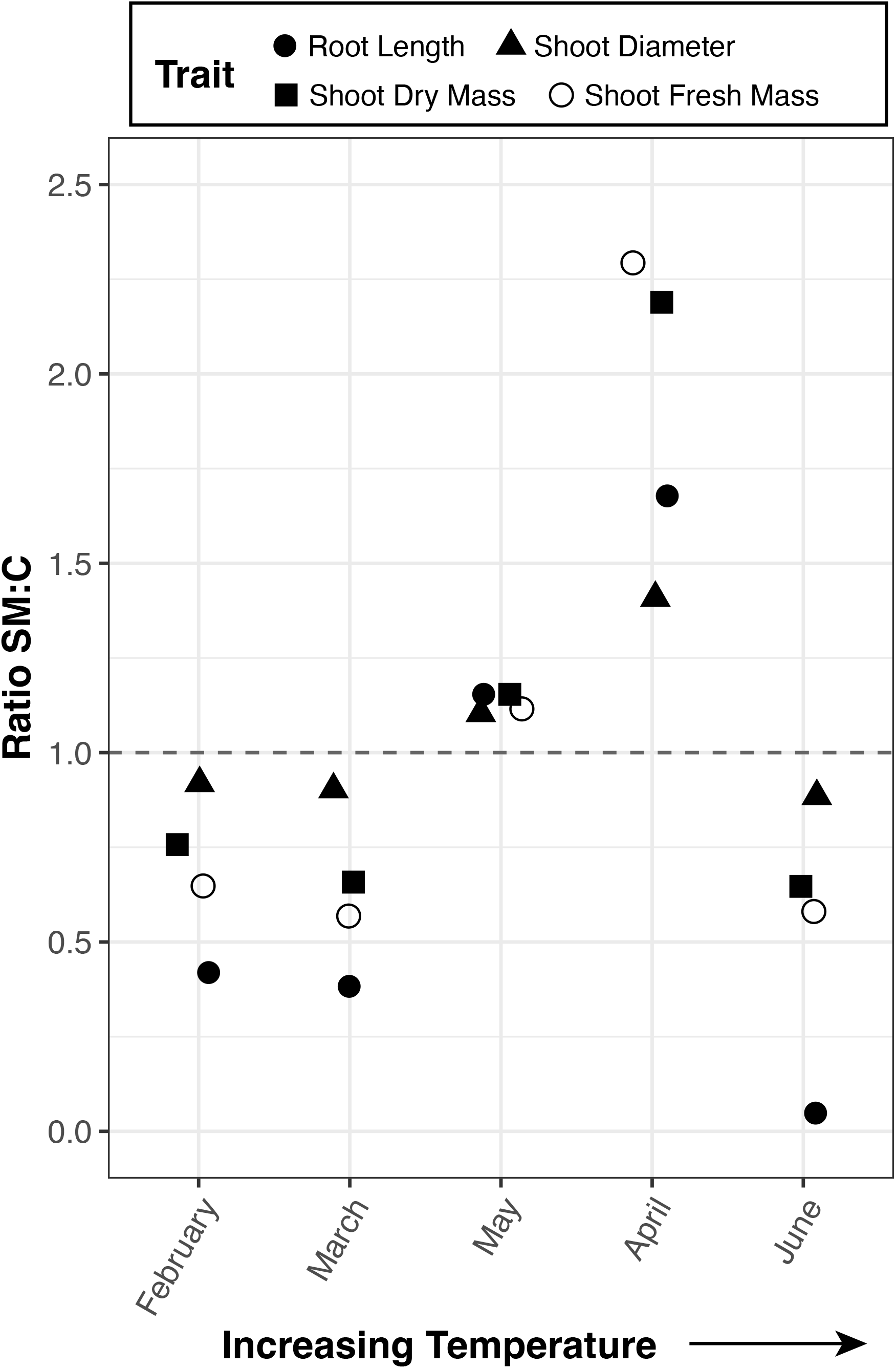
Plant traits were impacted by trial, with the average trial temperature mirroring the plant response in control conditions. (A) In control conditions, plants from February, March, and May were overall larger above and belowground compared to plants from April and June (*P*<0.05), regardless of cultivar. When the average temperature within each trial was compared with plant traits, plants with larger above and belowground traits, were also exposed to lower average temperatures. (B) In simulated microgravity conditions, plants from February, March, and April were more similar above and belowground (*P*<0.05), regardless of cultivar. Plants from May were on average larger above and belowground, while plants from June were on average smaller above and belowground. These findings did not reflect the average trial temperature. (A-B) Black circles indicate outliers. Black stars indicate the average trial temperature. Trials that share a letter were not significantly different from one another within each plant trait and gravity condition (*P*>0.05).

In contrast to the upright control conditions, plants grown under simulated microgravity did not follow the same response groupings (**Figure 5B, Table S11-Table S12**). However, when we analyze the relative growth between control and simulated microgravity conditions, distinct response patterns emerge (**Figure 6**). Specifically, with moderate heat stress in May and April, the simulated microgravity plants are larger than the control. For May, shoot traits under simulated microgravity are equivalent to or even exceed that of the February and March control plants under standard temperature conditions (**Figure 4, Figure 6**). These results suggest that moderate heat stress can alleviate the negative impacts of simulated microgravity on plant growth.

**Figure 6.**
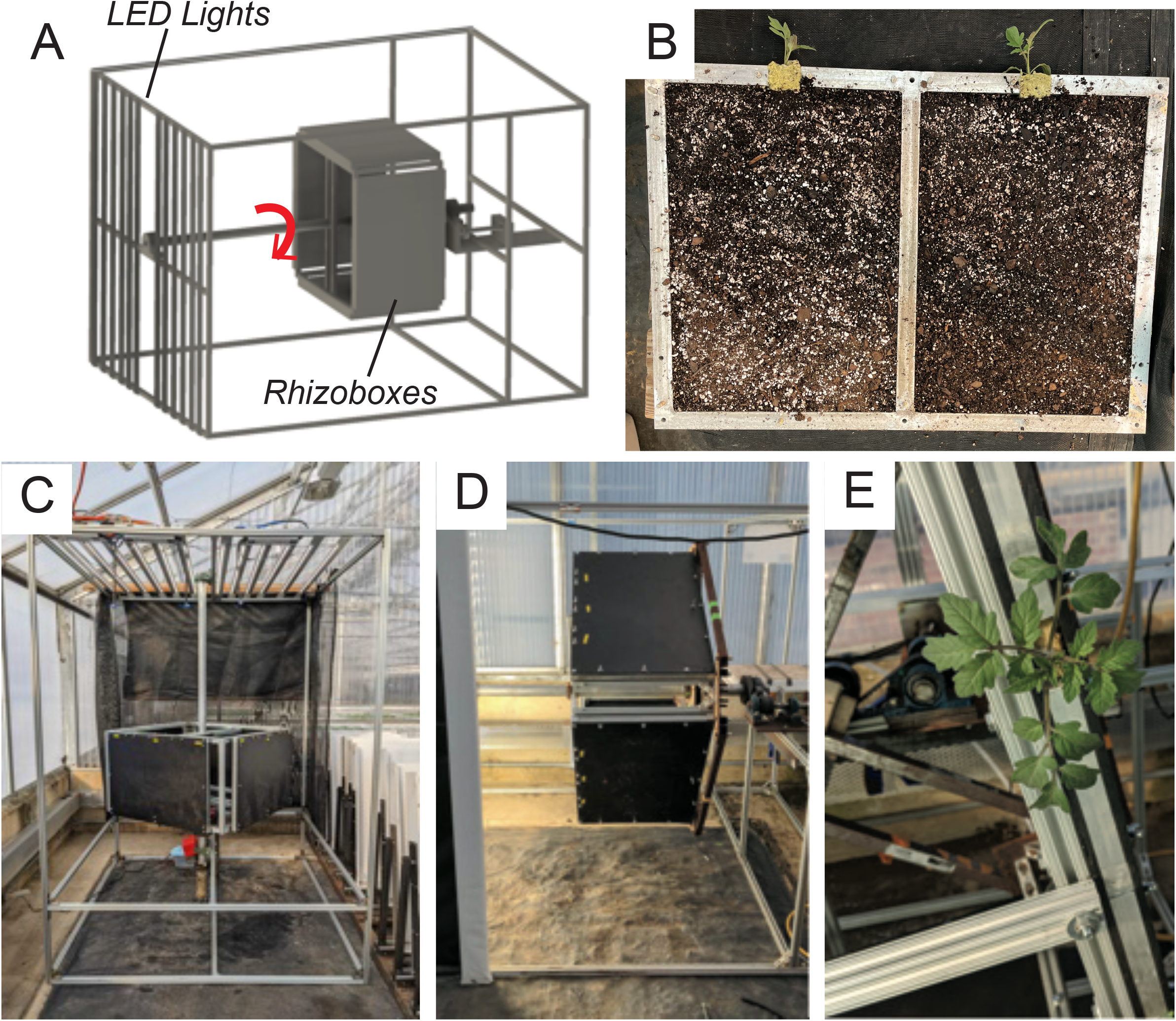
Trial-dependent trait responses to simulated microgravity. Within each trial, the average trait value in simulated microgravity conditions was compared to the average trait value in control conditions. For February, March, and June all traits were reduced in response to simulated microgravity, while in May and April all traits were enhanced in response to simulated microgravity.

## 4. Discussion and Conclusions

Understanding how plants alter their growth in response to microgravity requires an integration of spaceflight and ground-based approaches. Ground-based microgravity simulators are one approach to isolate the effects of gravity but are often limited in size and scope (Hasenstein and Loon, 2015). This study reports a set of meter-scale 2D clinostats that support plant growth beyond the seedling stage and expands the capacity to study gravitational impacts on plant growth and development. Using this system, we showed an environmentally dependent response to simulated microgravity.

A caveat of ground-based microgravity simulators pertains to their approach to randomize gravity through rotation. Rotation inherently introduces mechanical stress, which can be difficult to deconvolve from gravity randomization. To overcome this, we elected to use a rotating control that would also introduce mechanical stress. It is possible that the responses we observed were due to mechanical stimulation instead of gravity randomization. However, the stereotypical response of plants to mechanical stimulation is shorter and thicker stems, which was not observed. The different plant responses by trial with consistent mechanical stimulation further suggest that these responses are the result of gravity randomization instead of mechanical stimulation.

Our results suggest that elevated temperature is responsible for differences in both control and simulated microgravity responses in tomatoes (*S. lycopersicum*). Specifically, that moderate heat stress can alleviate the negative impacts of simulated microgravity. Whether this is a common attribute for many plant species or is specific to tomatoes is not clear.

The idea of one stressor alleviating another has been previously identified. For example, moderate heat stress has been shown to alleviate waterlogging stress in tomato (Wen et al., 2024). In the context of microgravity, root system impacts can be alleviated by providing light (David et al., 2025). However, the coordination between different stressors has been poorly explored and to our knowledge, this is the first data that suggests moderate heat stress may alleviate the detrimental impacts of plant growth for growing plants in simulated microgravity.

These data further highlight the trade-offs associated with exploring mature plant responses to simulated microgravity. The sheer size of the clinostats prevents them from inhabiting most growth chambers, and greenhouses are only semi-controlled environments. For the best environmental control in a greenhouse, trials would need to occur in the same month each year, but that would impose significant delays in research results and is not a feasible experimental design. Thus, future experiments should consider a scaled down the design to fit within a growth chamber that would enable testing of specific environmental interactions.

Collectively, this work demonstrates the utility of using large-scale 2D clinostats to study crop growth and development beyond the seedling stage and identifies an environmental factor that may alleviate the detrimental impacts of plant growth in simulated microgravity. Future experiments will focus on elucidating the relationship between temperature and microgravity.

## Abbreviations

ANOVA: analysis of variance
H7996: Hawaii7996
LED: Light-Emitting Diode
MM: MoneyMaker
PCA: Principal Component Analysis
PC: Principal Component
RPM: revolutions per minute

## Availability of Data and Materials

All raw data, processed data, and scripts used to analyze data are available on GitHub: https://github.com/EESparksLab/Hostetler_Kennebeck_et_al_2026

## CRediT authorship contribution statement

**Ashley N. Hostetler:** Formal analysis, Visualization, Writing – Original draft, Writing – Review & Editing

**Emily Kennebeck:** Investigation, Methodology, Project administration, Writing – Review & Editing

**Jonathan W. Reneau:** Investigation, Methodology, Visualization, Writing – Review & Editing

**Eva Birtell:** Investigation, Methodology, Writing – Review & Editing

**Denise L. Caldwell:** Investigation, Writing – Review & Editing

**Anjali S. Iyer-Pascuzzi:** Conceptualization, Funding acquisition, Methodology, Project administration, Supervision, Writing – Review & Editing

**Erin E. Sparks:** Conceptualization, Funding acquisition, Methodology, Project administration, Supervision, Visualization, Writing – Original draft, Writing – Review & Editing

## Acknowledgements

We gratefully acknowledge members of the Sparks Lab who assisted with clinostat set-up and take-down, specifically Dave Griffin and Austin Jensen. We also acknowledge the members of the University of Delaware senior design team (Emma Richmond-Boudewyns, Rachel Dennin, Peter Moreau, Paolo Tiamson, and Michaella Becker) who built the first prototype upon which the clinostats were designed.

## Supplementary Figure and Table Legends

**Figure S1. PCA of plant traits shows overlap by *Fusarium oxysporum* inoculation and mock inoculation**. (A) A PCA on plants in control and simulated microgravity conditions shows that PC1 and PC2 explain approximately 91% of the phenotypic variance. (B) A PCA on plants in upright control conditions shows that PC1 and PC2 explain approximately 94% of the phenotypic variance. (C) A PCA on plants in simulated microgravity shows that PC1 and PC2 explain approximately 93% of the phenotypic variance. (A-C) Each point represents a single plant across one of the five trials. There was complete overlap in phenotypic profiles for plants inoculated with *F. oxysporum* and those with mock inoculation.

**Figure S2. Example root images for two rhizoboxes**. An example of plant data that was excluded from analysis due to soil disruption.

**Figure S3. PCA of plant traits by trial reveals separation by cultivar**. Within each trial, a PCA of plant traits shows clear separation by cultivar along PC2. PC2 explains 11-25% of the phenotypic variation and is dependent on trial.

**Figure S4. Temperature profiles varied across trials. (**A) The average daily temperature across trials was consistent within a trial except for June, which had a temperature spike from days 7-12. (B) The average hourly temperature across trials showed increasing values throughout the day across trials.

**Figure S5. Apical bud height is dependent on tomato cultivar**. Regardless of gravity condition or trial, cultivar Moneymaker is on average shorter than cultivar Hawaii7996. Within each trial, cultivars that share a letter are not significantly different than one another (*P*>0.05). Black dots indicate outliers.

**Figure S6. Percent dry mass varied by trial**. The percent dry mass is a measure of the total plant mass after drying and is a good indicator of plant water status. Regardless of cultivar, the impact of gravity condition was variable and impacted by trial (*P*>0.05). Gravity conditions that share a letter are not significantly different from one another within each trial (*P*>0.05). Black circles indicate outliers.

**Figure S7. Plant traits were impacted by trial, with the accumulation of solar radiation mirroring the plant response in control conditions**. (A) In control conditions, plants from February, March, and May were overall larger above and belowground compared to plants from April and June (*P*<0.05), regardless of cultivar. When the cumulative solar radiation within each trial was compared with plant traits, plants with larger above and belowground traits were also exposed to lower amounts of cumulative solar radiation. (B) In simulated microgravity conditions, plants from February, March, and April were more similar above and belowground (*P*<0.05), regardless of cultivar. Plants from May were on average larger above and belowground, while plants from June were on average smaller above and belowground. These findings did not reflect the cumulative solar radiation within each trial. (A-B) Black circles indicate outliers. Black triangles indicate the amount of cumulative solar radiation within each trial. Trials that share a letter were not significantly different from one another within each plant trait and gravity condition (*P*>0.05).

**Table S1. Parts List**. A List of all the parts required to construct one 2D clinostat for supporting plant growth beyond the seedling stage.

**Table S2. Phenotypic traits and their eigenvector loadings from a PCA on all trials**. A principal component analysis (PCA) was performed on the following traits: apical bud height, shoot fresh mass, shoot dry mass, shoot diameter, and root length. The eigenvector loadings for the first two principal components (PC1 and PC2) were extracted for each of the traits.

**Table S3. Phenotypic traits and their eigenvector loadings for each trial for PC1**. Individual principal component analyses were performed for each trial on the following traits: apical bud height, shoot fresh mass, shoot dry mass, shoot diameter, and root length. The eigenvector loadings for PC1 were extracted for each trial.

**Table S4. Phenotypic traits and their eigenvector loadings for each trial for PC2**. Individual principal component analyses were performed for each trial on the following traits: apical bud height, shoot fresh mass, shoot dry mass, shoot diameter, and root length. The eigenvector loadings for PC2 were extracted for each trial.

**Table S5. Environmental Data for each trial**. For each trial the environmental data used as covariates in the environmental model are listed.

**Table S6. Likelihood ratio test between a base model and an environmental model for all trials**. The base model (trait ∼ Cultivar * Gravity Condition + (1|Trial) was compared to an environmental model to determine if including environmental factors as covariates reduce the variation due to trial. Six environmental models were tested: (1) (trait ∼ Cultivar * Gravity Condition + Temperature + Relative Humidity + Solar Radiation + (1| Trial)); (2) (trait ∼ Cultivar * Gravity Condition + Temperature + (1| Trial)); (3) (trait ∼ Cultivar * Gravity Condition Relative Humidity + (1| Trial)); (4) (trait ∼ Cultivar * Gravity Condition + Solar Radiation + (1| Trial)); (5) (trait ∼ Cultivar * Gravity Condition + Daytime Temp + (1| Trial)); (6) (trait ∼ Cultivar * Gravity Condition + Nighttime Temp + (1| Trial)).

**Table S7. ANOVA table for each plant trait within each trial**. To determine if significant differences were observed between cultivar, gravity condition, or their interaction, a two-way analysis of variance (ANOVA) was used for each trait within each trial. Within each model, cultivar and gravity condition were included as independent variables. A Tukey’s ladder of powers transformation was used if residuals were not from a normal distribution.

**Table S8. Pairwise comparison summary table for each plant trait within each trial**. For each plant trait and each trial, the estimated marginal means, standard error, degrees of freedom, lower confidence level, upper confidence level, and Tukey HSD groupings (Group) are shown for each main effect (gravity condition or cultivar).

**Table S9. ANOVA table for each plant trait within upright control conditions**. To determine if significant differences were observed between cultivar or trial, a two-way analysis of variance (ANOVA) was used for each trait within upright control conditions. Within each model, cultivar and trial were included as independent variables.

**Table S10. Pairwise comparison summary table for each plant trait in upright control conditions**. For each plant trait, summary statistics and Tukey HSD groupings (Group) are shown for each trial in upright control conditions.

**Table S11. ANOVA table for each plant trait within simulated microgravity conditions**. To determine if significant differences were observed between cultivar or trial, a two-way analysis of variance (ANOVA) was used for each trait within simulated microgravity conditions. Within each model, cultivar and trial were included as independent variables. A Tukey’s ladder of powers transformation was used if residuals were not from a normal distribution.

**Table S12. Pairwise comparison summary table for each plant trait in simulated microgravity conditions**. For each plant trait, summary statistics and Tukey HSD groupings (Group) are shown for each trial in simulated microgravity conditions.

