## Supplementary figures and images for "Meter-scale 2D clinostats uncover environmentally derived variation in tomato responses to simulated microgravity"

### FIgure S1

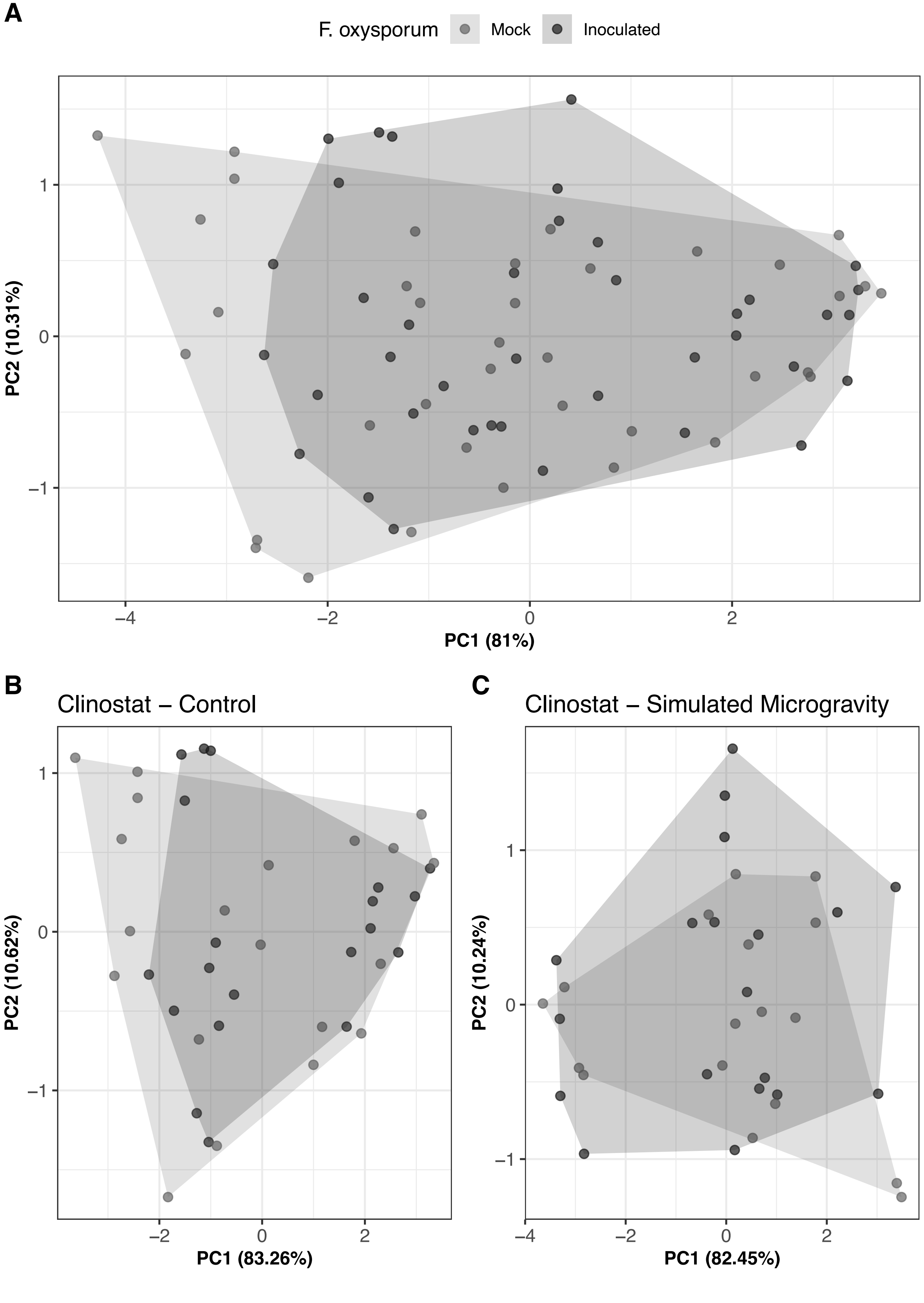

### Figure S2

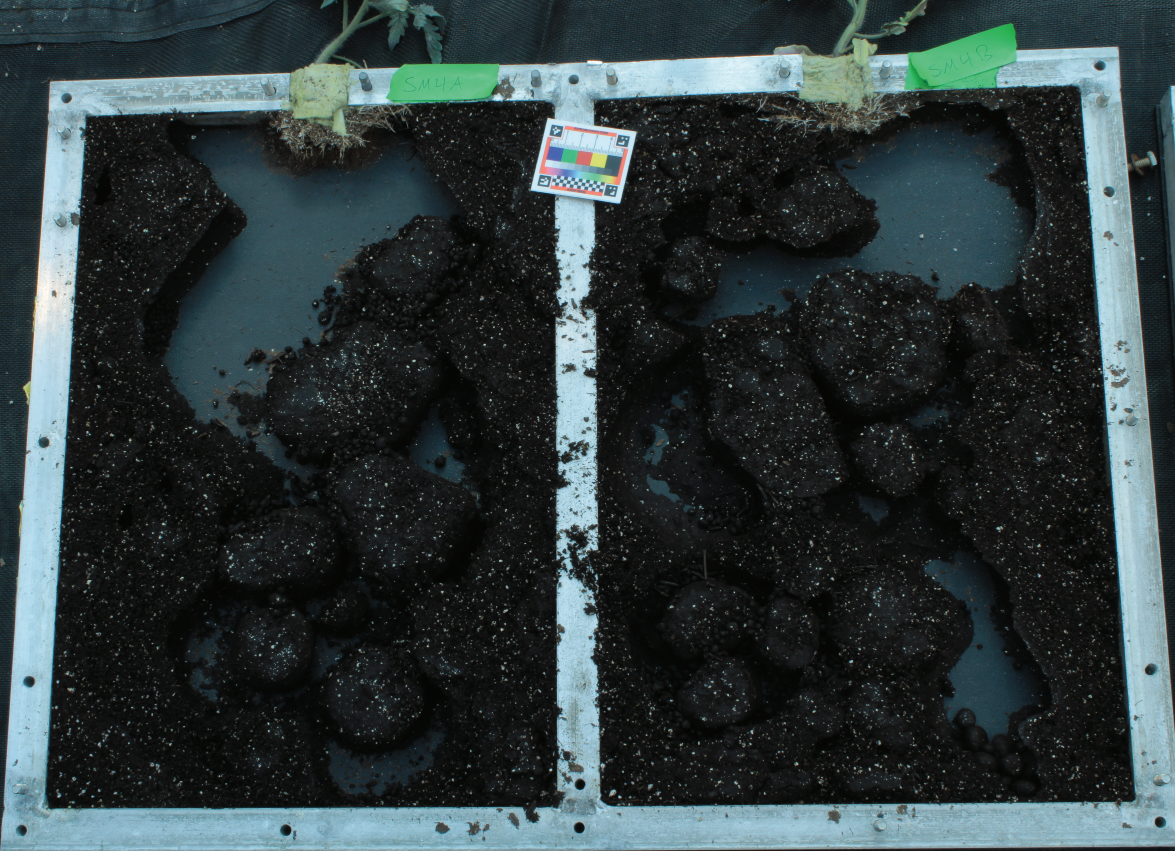

### FIgure S3

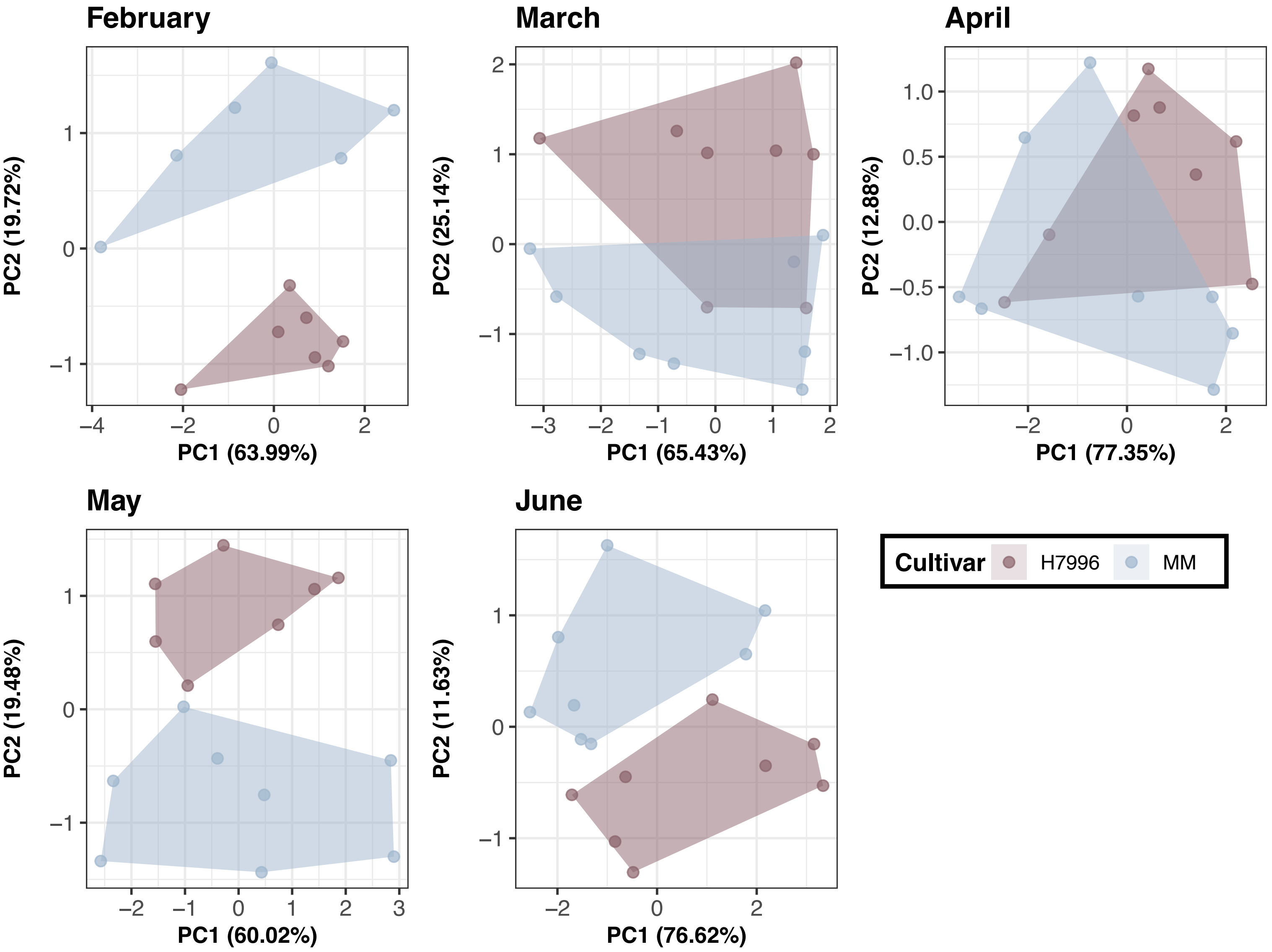

### FIgure S4

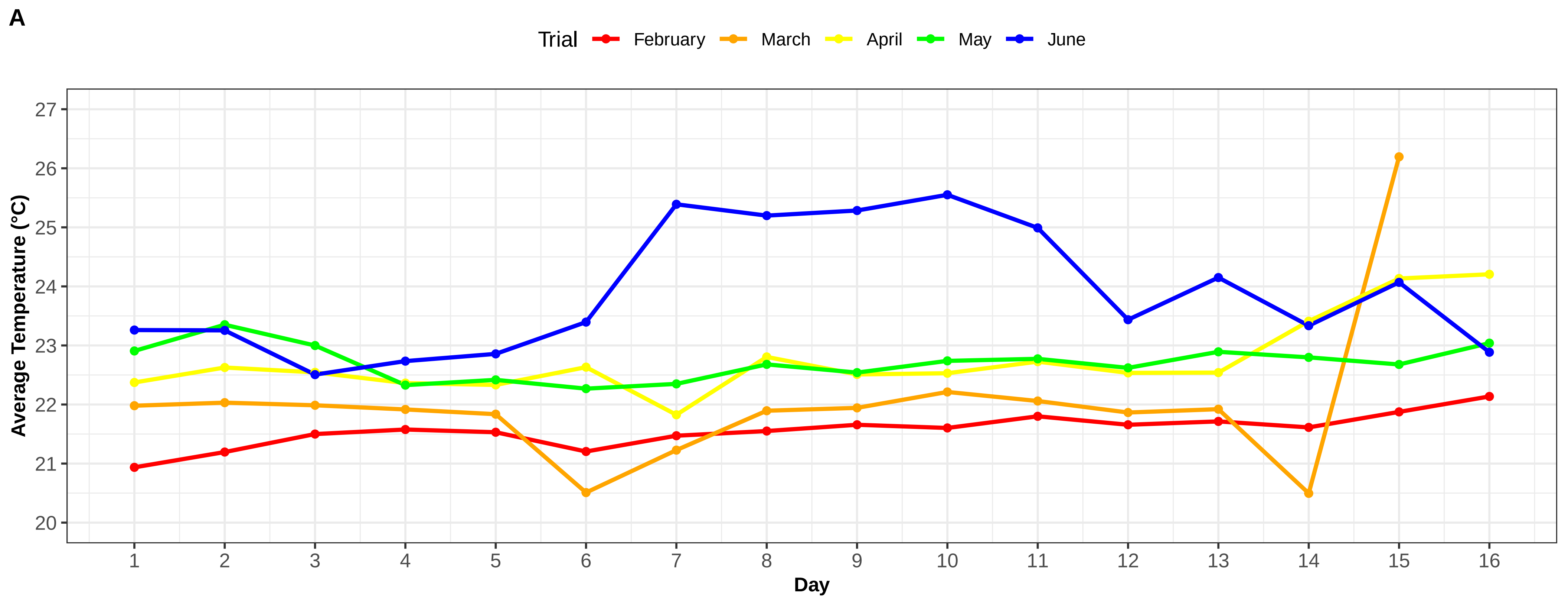

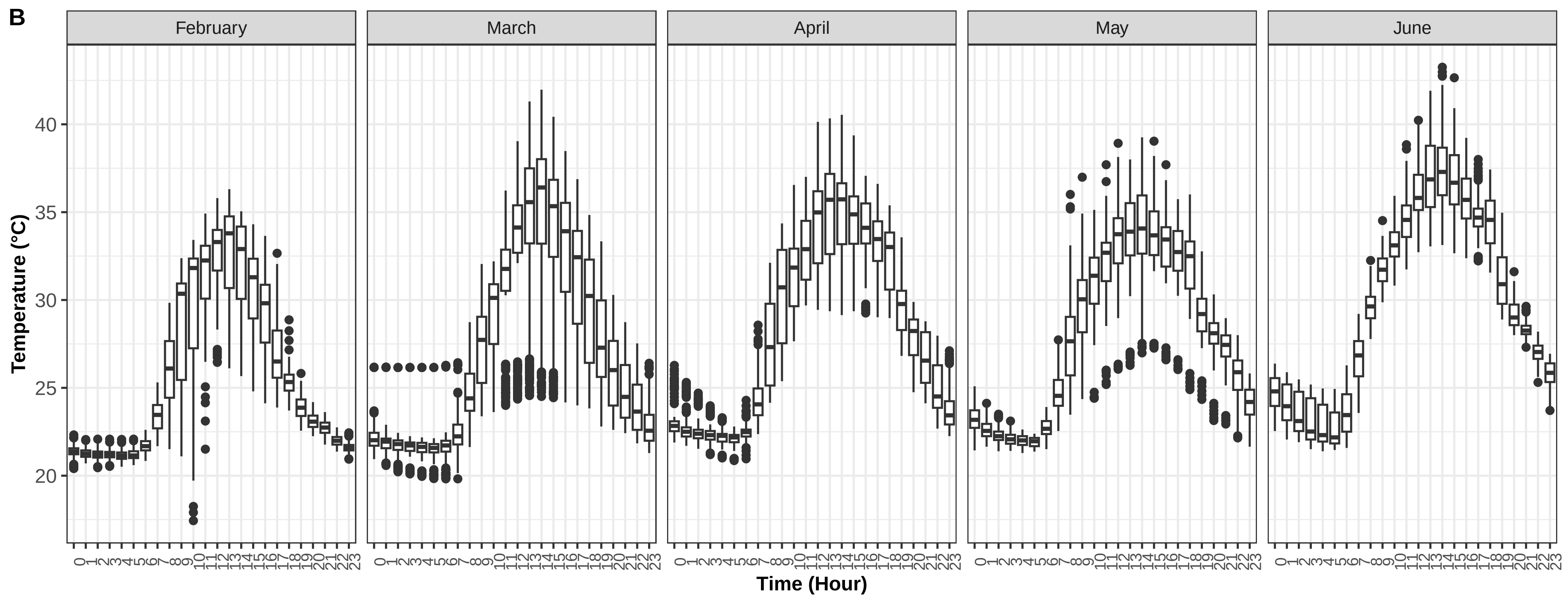

### FIgure S5

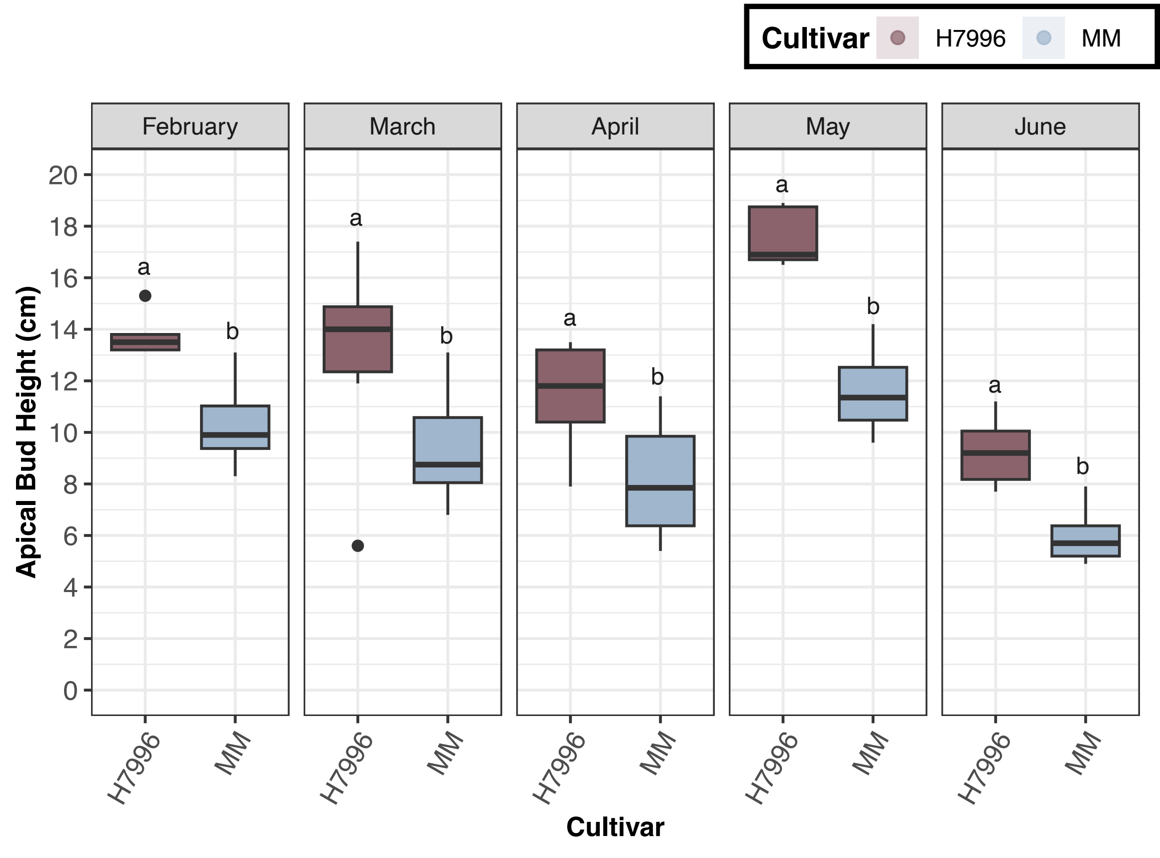

### FIgure S6

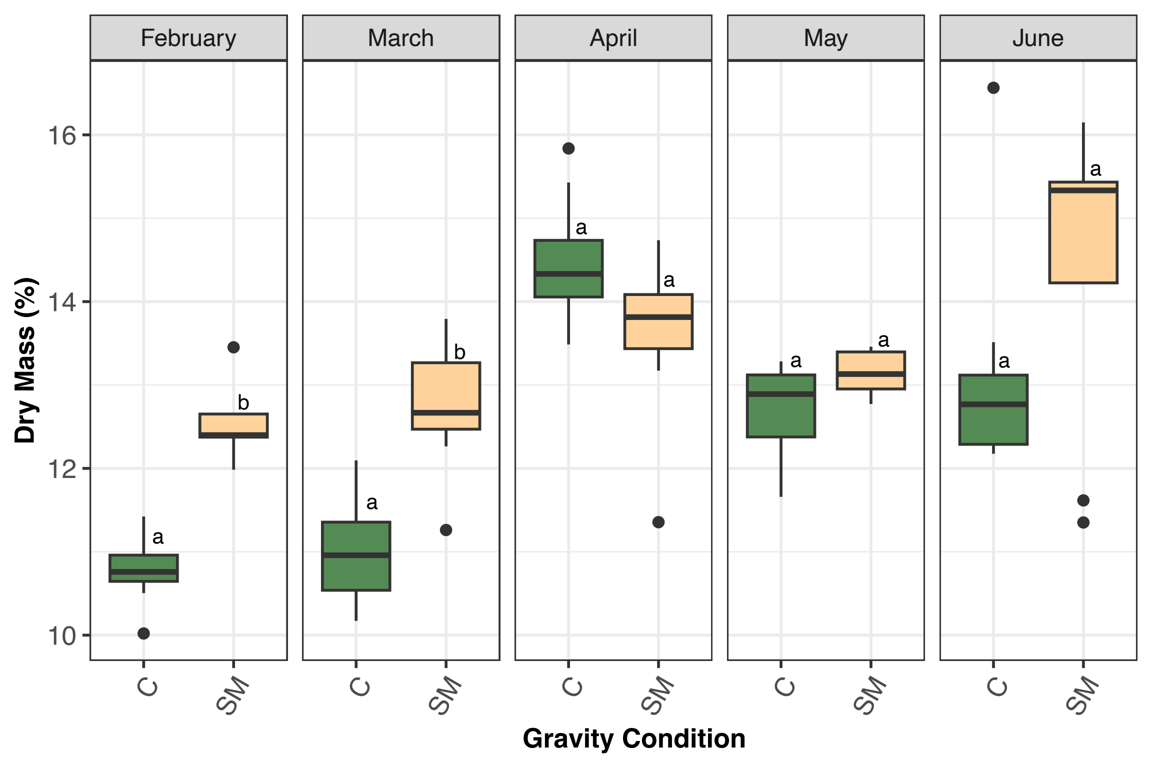

### FIgure S7

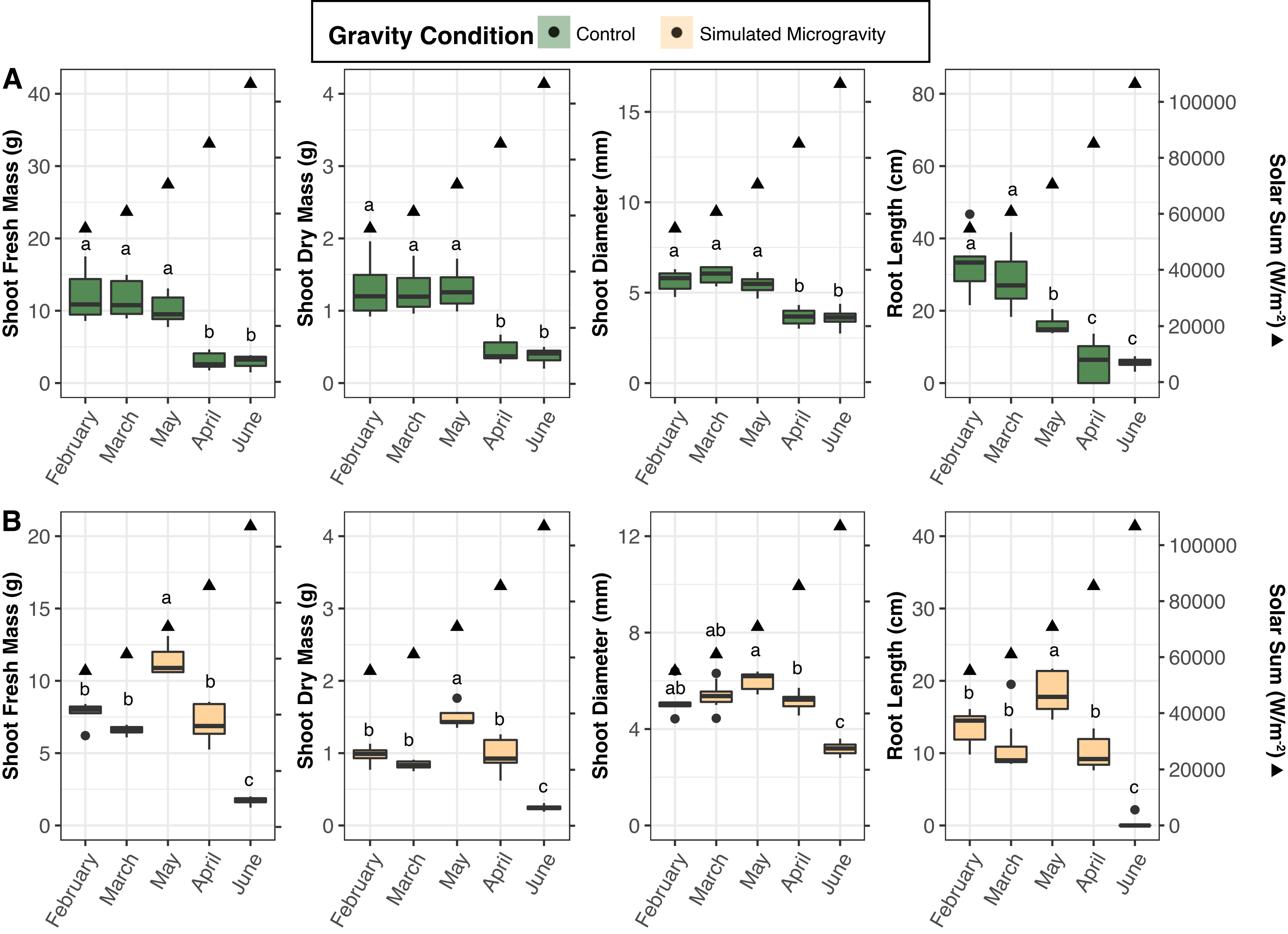
