## Supplemental Tables for "Meter-scale 2D clinostats uncover environmentally derived variation in tomato responses to simulated microgravity"

**Table S1. Parts List.** List of all the parts required to construct one 2D clinostat for supporting plant growth beyond the seedling stage.

| Number | Vendor | Part Number | Description |
|---|---|---|---|
| 1 | Automation Direct | STL40L100AF-2012 | SureMotion timing pulley, ductile iron, 3/8in L pitch, 40 tooth, 4.775in pitch diameter, no hub with flanges. For use with 1in wide belt. Requires 2012 series Taper-Lock style bushing |
| 1 | Automation Direct | QD-SH-1000 | SureMotion QD bushing, SH size, 1.000in bore, steel |
| 1 | Automation Direct | SQD24L100AF-SH | (SureMotion timing pulley, steel, 3/8in L pitch, 24 tooth, 2.865in pitch diameter, no hub with flanges. For use with 1in wide belt. Requires SH series QD style bushing.) |
| 1 | makermotor.com | PN00113-VAR | - Variable Speed High Torque 5 RPM Conveyor and Rotisserie Gear Motor 12V DC Reversible |
| 1 | makermotor.com | PN00409 | 12v Gear Motor PWM Variable Speed 12vdc Gearmotor |
| 1 | McMaster Carr | 9056K14 | Multipurpose 6061 Aluminum Round Tube, 3/8" Wall Thickness, 2" OD, 6 Feet Long, |
| 4 | McMaster Carr | 47065T101 | T-Slotted Framing, Single 4-Slot Rail, Silver, 1" High x 1" Wide, Solid, 7' Long, |
| 12 | McMaster Carr | 47065T101 | T-Slotted Framing, Single 4-Slot Rail, Silver, 1" High x 1" Wide, Solid, 5' Long, |
| 12 | Heatsink USA | | 2.079" WIDE EXTRUDED ALUMINUM HEATSINK, 62" Long |
| 2 | Amazon | PGN - UCP210-32 | Pillow Block Mounted Ball Bearing 2" Bore Self Aligning |
| 4 | Amazon | Climax Metal 2C-200 | Steel Two-Piece Clamping Collar, Black Oxide Plating, 2" Bore Size, 3" OD, With 5/16-24 x 1 Set Screw |
| 1 | Amazon | Timken T101 904A1 | Tapered Roller Thrust Bearing, Chromium Steel |
| 4 | US Plastics | | 1/4" x 48" x 48" Black Seaboard¬Æ UV Stabilized HDPE Sheet |
| 1 | Steel and Metal Service Center | | 2"x2"x10'angle steel stock to make custom 2"x2"x1" corner brackets |
| 200 | Albany County Fasteners | 15002 | Button Socket Head Screws 304 SS - 1/4-20 x 3/8 |
| 24 | Albany County Fasteners | 15020 | Button Socket Head Screws 304 SS - 1/4-20 x 3/4 |
| 1 | Diesel Belting | 480L100 | Timing Belt 0.375in Pitch, 1in Wide, 48in Pitch Length, 128 Teeth |
| 1 | Amazon | ACDelco Gold 38159 | Drive Belt Tensioner Assembly with Pulley |

**Table S2. Phenotypic traits and their eigenvector loadings from a PCA on all trials.** A principal component analysis (PCA) was performed on the following traits: apical bud height, shoot fresh mass, shoot dry mass, shoot diameter, and root length. The eigenvector loadings for the first two principal components (PC1 and PC2) were extracted for each of the traits.

| | PC1 | PC2 |
|---|---|---|
| **Apical Bud Height** | -0.350 | -0.882 |
| **Shoot Fresh Mass** | -0.509 | 0.106 |
| **Shoot Dry Mass** | -0.496 | -0.002 |
| **Shoot Diameter** | -0.442 | 0.175 |
| **Root Length** | -0.421 | 0.424 |

**Table S3. Phenotypic traits and their eigenvector loadings for each trial for PC1.** Individual principal component analyses were performed for each trial on the following traits: apical bud height, shoot fresh mass, shoot dry mass, shoot diameter, and root length. The eigenvector loadings for PC1 were extracted for each trial.

| | February | March | April | May | June |
|---|---|---|---|---|---|
| **Apical Bud Height** | -0.170 | -0.297 | 0.411 | -0.181 | 0.400 |
| **Shoot Fresh Mass** | -0.550 | -0.548 | 0.551 | -0.551 | 0.508 |
| **Shoot Dry Mass** | -0.538 | -0.531 | 0.538 | -0.547 | 0.501 |
| **Shoot Diameter** | -0.394 | -0.238 | 0.477 | -0.512 | 0.482 |
| **Root Length** | -0.473 | -0.522 | 0.104 | -0.320 | 0.315 |

**Table S4. Phenotypic traits and their eigenvector loadings for each trial for PC2.** Individual principal component analyses were performed for each trial on the following traits: apical bud height, shoot fresh mass, shoot dry mass, shoot diameter, and root length. The eigenvector loadings for PC2 were extracted for each trial.

| | February | March | April | May | June |
|---|---|---|---|---|---|
| **Apical Bud Height** | -0.949 | 0.670 | 0.408 | 0.924 | -0.356 |
| **Shoot Fresh Mass** | -0.021 | -0.025 | -0.091 | -0.028 | -0.064 |
| **Shoot Dry Mass** | -0.024 | 0.005 | -0.124 | -0.154 | 0.067 |
| **Shoot Diameter** | 0.257 | -0.741 | -0.293 | 0.079 | -0.286 |
| **Root Length** | 0.179 | -0.022 | 0.851 | -0.339 | 0.885 |

**Table S5.** Environmental Data for each trial. For each trial the environmental data used as covariates in the environmental model are listed.

| Trial | Cumulative Solar Radiation | Average Temperature (degC) | SD Temperature (degC) | Average Relative Humidity (%) | SD Relative Humidity (%) |
|---|---|---|---|---|---|
| February | 55153.00 | 25.47 | 4.53 | 29.71 | 13.32 |
| March | 61108.00 | 26.78 | 5.22 | 37.05 | 18.11 |
| April | 85464.00 | 28.18 | 5.18 | 37.92 | 15.59 |
| May | 70875.00 | 27.77 | 4.78 | 49.65 | 15.46 |
| June | 106795.00 | 29.70 | 5.36 | 44.57 | 19.70 |

**Table S6.** Likelihood ratio test between a base model and an environmental model for all trials. The base model (trait ~ Cultivar * Gravity Condition + (1|Trial) was compared to an environmental model to determine if including environmental factors as covariates reduce the variation due to trial. Six environmental models were tested: (1) (trait ~ Cultivar * Gravity Condition + Temperature + Relative Humidity + Solar Radiation + (1| Trial)); (2) (trait ~ Cultivar * Gravity Condition + Temperature + (1| Trial)); (3) (trait ~ Cultivar * Gravity Condition Relative Humidity + (1| Trial)); (4) (trait ~ Cultivar * Gravity Condition + Solar Radiation + (1| Trial)); (5) (trait ~ Cultivar * Gravity Condition + Daytime Temp + (1| Trial)); (6) (trait ~ Cultivar * Gravity Condition + Nighttime Temp + (1| Trial)).

| | **Chisq (Base vs Env)** | ***P* (Base vs Env)** | **Replication SD (base)** | **Residual SD (base)** | **Replication SD (Env)** | **Residual SD (Env)** |
|---|---|---|---|---|---|---|
| Apical Bud Height | 15.37 | 0.002 | 2.23 | 1.94 | 0.07 | 1.94 |
| Shoot Fresh Mass | 21.85 | 0.000 | 3.20 | 2.35 | 0.00 | 2.27 |
| Shoot Dry Mass | 18.57 | 0.000 | 0.37 | 0.27 | 0.00 | 0.26 |
| Shoot Diameter | 22.15 | 0.000 | 0.89 | 0.63 | 0.00 | 0.61 |
| Root Length | 18.81 | 0.000 | 7.58 | 6.50 | 0.00 | 6.36 |

| | **Chisq (Base vs Temp)** | ***P* (Base vs Temp)** | **Replication SD (base)** | **Residual SD (base)** | **Replication SD (Temp)** | **Residual SD (Temp)** |
|---|---|---|---|---|---|---|
| Apical Bud Height | 2.05 | 0.152 | 2.23 | 1.94 | 1.79 | 1.94 |
| Shoot Fresh Mass | 5.52 | 0.019 | 3.20 | 2.35 | 1.77 | 2.35 |
| Shoot Dry Mass | 3.82 | 0.051 | 0.37 | 0.27 | 0.25 | 0.27 |
| Shoot Diameter | 5.53 | 0.019 | 0.89 | 0.63 | 0.49 | 0.63 |
| Root Length | 12.25 | 0.000 | 7.58 | 6.50 | 1.56 | 6.50 |

| | **Chisq (Base vs RH)** | ***P* (Base vs RH)** | **Replication SD (base)** | **Residual SD (base)** | **Replication SD (RH)** | **Residual SD (RH)** |
|---|---|---|---|---|---|---|
| Apical Bud Height | 0.10 | 0.752 | 2.23 | 1.94 | 2.21 | 1.94 |
| Shoot Fresh Mass | 0.16 | 0.688 | 3.20 | 2.35 | 3.14 | 2.35 |
| Shoot Dry Mass | 0.01 | 0.907 | 0.37 | 0.27 | 0.37 | 0.27 |
| Shoot Diameter | 0.24 | 0.625 | 0.89 | 0.63 | 0.87 | 0.63 |
| Root Length | 1.38 | 0.240 | 7.58 | 6.50 | 6.56 | 6.50 |

| | **Chisq (Base vs Solar)** | ***P* (Base vs Solar)** | **Replication SD (base)** | **Residual SD (base)** | **Replication SD (Solar)** | **Residual SD (Solar)** |
|---|---|---|---|---|---|---|
| Apical Bud Height | 3.95 | 0.047 | 2.23 | 1.94 | 1.46 | 1.94 |
| Shoot Fresh Mass | 9.61 | 0.002 | 3.20 | 2.35 | 1.09 | 2.35 |
| Shoot Dry Mass | 6.82 | 0.009 | 0.37 | 0.27 | 0.18 | 0.27 |
| Shoot Diameter | 11.09 | 0.001 | 0.89 | 0.63 | 0.25 | 0.63 |
| Root Length | 16.63 | 0.000 | 7.58 | 6.50 | 0.00 | 6.45 |

**DAYTIME ONLY**

| | **Chisq (Base vs Temp)** | ***P* (Base vs Temp)** | **Replication SD (base)** | **Residual SD (base)** | **Replication SD (Temp)** | **Residual SD (Temp)** |
|---|---|---|---|---|---|---|
| Apical Bud Height | 1.90 | 0.168 | 2.23 | 1.94 | 1.82 | 1.94 |
| Shoot Fresh Mass | 5.28 | 0.022 | 3.20 | 2.35 | 1.82 | 2.35 |
| Shoot Dry Mass | 3.61 | 0.058 | 0.37 | 0.27 | 0.25 | 0.27 |
| Shoot Diameter | 5.36 | 0.021 | 0.89 | 0.63 | 0.50 | 0.63 |
| Root Length | 12.09 | 0.001 | 7.58 | 6.50 | 1.61 | 6.50 |

**NIGHTTIME ONLY**

| | **Chisq (Base vs Temp)** | ***P* (Base vs Temp)** | **Replication SD (base)** | **Residual SD (base)** | **Replication SD (Temp)** | **Residual SD (Temp)** |
|---|---|---|---|---|---|---|
| Apical Bud Height | 2.07 | 0.150 | 2.23 | 1.94 | 1.79 | 1.94 |
| Shoot Fresh Mass | 5.36 | 0.021 | 3.20 | 2.35 | 1.80 | 2.35 |
| Shoot Dry Mass | 3.88 | 0.049 | 0.37 | 0.27 | 0.25 | 0.27 |
| Shoot Diameter | 6.20 | 0.013 | 0.89 | 0.63 | 0.46 | 0.63 |
| Root Length | 10.33 | 0.001 | 7.58 | 6.50 | 2.22 | 6.50 |

**Table S7. ANOVA table for each plant trait within each trial.** To determine if significant differences were observed between cultivar, gravity condition, or their interaction, a two-way analysis of variance (ANOVA) was used for each trait within each trial. Within each model, cultivar and gravity condition were included as independent variables. A Tukey's ladder of powers transformation was used if residuals were not from a normal distribution.

| Trait | Trial | | Estimate | Std. Erro | t value | Pr(>\|t\|) | Transformation |
|---|---|---|---|---|---|---|---|
| Apical Bud Height | 1 | (Intercept) | 13.95 | 0.57 | 24.45 | 1.54E-09 | None |
| | | Cultivar (MM) | -2.98 | 0.81 | -3.69 | 5.02E-03 | |
| | | Clinostat (SM) | -0.55 | 0.87 | -0.63 | 5.44E-01 | |
| | | Cultivar (MM)*Clinostat (SM) | -1.48 | 1.32 | -1.12 | 2.92E-01 | |
| | 2 | (Intercept) | 15.05 | 1.21 | 12.41 | 3.32E-08 | |
| | | Cultivar (MM) | -4.00 | 1.71 | -2.33 | 3.79E-02 | |
| | | Clinostat (SM) | -3.68 | 1.71 | -2.14 | 5.33E-02 | |
| | | Cultivar (MM)*Clinostat (SM) | 0.35 | 2.43 | 0.14 | 8.88E-01 | |
| | 3 | (Intercept) | 10.30 | 0.83 | 12.35 | 3.51E-08 | |
| | | Cultivar (MM) | -3.93 | 1.18 | -3.33 | 6.02E-03 | |
| | | Clinostat (SM) | 2.15 | 1.18 | 1.82 | 9.33E-02 | |
| | | Cultivar (MM)*Clinostat (SM) | 1.55 | 1.67 | 0.93 | 3.71E-01 | |
| | 4 | (Intercept) | 17.20 | 0.73 | 23.40 | 9.84E-11 | |
| | | Cultivar (MM) | -4.88 | 1.04 | -4.69 | 6.60E-04 | |
| | | Clinostat (SM) | 0.93 | 1.12 | 0.83 | 4.23E-01 | |
| | | Cultivar (MM)*Clinostat (SM) | -2.31 | 1.53 | -1.51 | 1.60E-01 | |
| | 5 | (Intercept) | 10.18 | 0.52 | 19.75 | 1.62E-10 | |
| | | Cultivar (MM) | -3.78 | 0.73 | -5.18 | 2.29E-04 | |
| | | Clinostat (SM) | -1.75 | 0.73 | -2.40 | 3.34E-02 | |
| | | Cultivar (MM)*Clinostat (SM) | 0.88 | 1.03 | 0.85 | 4.12E-01 | |
| **Trait** | **Trial** | | **Estimate** | **Std. Erro** | **t value** | **Pr(>\|t\|)** | **Transformation** |
| Shoot Fresh Mass | 1 | (Intercept) | 10.61 | 1.32 | 8.06 | 0.00 | None |
| | | Cultivar (MM) | 2.62 | 1.86 | 1.41 | 0.19 | |
| | | Clinostat (SM) | -2.55 | 2.01 | -1.27 | 0.24 | |
| | | Cultivar (MM)*Clinostat (SM) | -3.45 | 3.04 | -1.14 | 0.28 | |
| | 2 | (Intercept) | 10.40 | 0.85 | 12.17 | 0.00 | |
| | | Cultivar (MM) | 2.37 | 1.21 | 1.96 | 0.07 | |
| | | Clinostat (SM) | -3.74 | 1.21 | -3.10 | 0.01 | |
| | | Cultivar (MM)*Clinostat (SM) | -2.51 | 1.71 | -1.47 | 0.17 | |
| | 3 | (Intercept) | 3.51 | 0.64 | 5.52 | 0.00 | |
| | | Cultivar (MM) | -0.87 | 0.90 | -0.96 | 0.35 | |
| | | Clinostat (SM) | 3.32 | 0.90 | 3.69 | 0.00 | |
| | | Cultivar (MM)*Clinostat (SM) | 1.32 | 1.27 | 1.04 | 0.32 | |
| | 4 | (Intercept) | 9.86 | 0.86 | 11.41 | 0.00 | |
| | | Cultivar (MM) | 0.71 | 1.22 | 0.58 | 0.57 | |
| | | Clinostat (SM) | 1.67 | 1.32 | 1.26 | 0.23 | |
| | | Cultivar (MM)*Clinostat (SM) | -0.95 | 1.80 | -0.53 | 0.61 | |
| | 5 | (Intercept) | 3.36 | 0.33 | 10.31 | 0.00 | |
| | | Cultivar (MM) | -0.84 | 0.46 | -1.83 | 0.09 | |
| | | Clinostat (SM) | -1.51 | 0.46 | -3.26 | 0.01 | |
| | | Cultivar (MM)*Clinostat (SM) | 0.54 | 0.65 | 0.83 | 0.42 | |
| **Trait** | **Trial** | | **Estimate** | **Std. Erro** | **t value** | **Pr(>\|t\|)** | **Transformation** |
| | 1 | (Intercept) | 1.13 | 0.15 | 7.77 | 0.00 | |
| | | Cultivar (MM) | 0.30 | 0.21 | 1.47 | 0.18 | |
| | | Clinostat (SM) | -0.12 | 0.22 | -0.52 | 0.62 | |
| | | Cultivar (MM)*Clinostat (SM) | -0.41 | 0.34 | -1.23 | 0.25 | |
| | 2 | (Intercept) | 1.13 | 0.09 | 12.03 | 0.00 | |
| | | Cultivar (MM) | 0.29 | 0.13 | 2.15 | 0.05 | |
| | | Clinostat (SM) | -0.30 | 0.13 | -2.22 | 0.05 | |
| | | Cultivar (MM)*Clinostat (SM) | -0.28 | 0.19 | -1.49 | 0.16 | |

| | | | | | | | |
|---|---|---|---|---|---|---|---|
| **Shoot Dry Mass** | 3 | (Intercept) | 0.51 | 0.10 | 4.97 | 0.00 | None |
| | | Cultivar (MM) | -0.14 | 0.15 | -0.93 | 0.37 | |
| | | Clinostat (SM) | 0.42 | 0.15 | 2.89 | 0.01 | |
| | | Cultivar (MM)*Clinostat (SM) | 0.21 | 0.21 | 1.04 | 0.32 | |
| | 4 | (Intercept) | 1.24 | 0.11 | 10.98 | 0.00 | |
| | | Cultivar (MM) | 0.11 | 0.16 | 0.70 | 0.50 | |
| | | Clinostat (SM) | 0.24 | 0.17 | 1.41 | 0.19 | |
| | | Cultivar (MM)*Clinostat (SM) | -0.09 | 0.24 | -0.39 | 0.70 | |
| | 5 | (Intercept) | 0.42 | 0.04 | 10.80 | 0.00 | |
| | | Cultivar (MM) | -0.08 | 0.06 | -1.50 | 0.16 | |
| | | Clinostat (SM) | -0.17 | 0.06 | -3.14 | 0.01 | |
| | | Cultivar (MM)*Clinostat (SM) | 0.08 | 0.08 | 1.00 | 0.34 | |
| **Trait** | **Trial** | | **Estimate** | **Std. Erro** | **t value** | **Pr(>\|t\|)** | **Transformation** |
| **Shoot Diameter** | 1 | (Intercept) | 5.17 | 0.22 | 23.15 | 0.00 | None |
| | | Cultivar (MM) | 0.94 | 0.32 | 2.96 | 0.02 | |
| | | Clinostat (SM) | 0.35 | 0.34 | 1.01 | 0.34 | |
| | | Cultivar (MM)*Clinostat (SM) | -1.77 | 0.52 | -3.43 | 0.01 | |
| | 2 | (Intercept) | 5.70 | 0.22 | 25.33 | 0.00 | |
| | | Cultivar (MM) | 0.55 | 0.32 | 1.73 | 0.11 | |
| | | Clinostat (SM) | -0.65 | 0.32 | -2.05 | 0.06 | |
| | | Cultivar (MM)*Clinostat (SM) | 0.14 | 0.45 | 0.32 | 0.76 | |
| | 3 | (Intercept) | 3.98 | 0.15 | 26.07 | 0.00 | |
| | | Cultivar (MM) | -0.65 | 0.22 | -2.99 | 0.01 | |
| | | Clinostat (SM) | 0.96 | 0.22 | 4.44 | 0.00 | |
| | | Cultivar (MM)*Clinostat (SM) | 1.07 | 0.31 | 3.51 | 0.00 | |
| | 4 | (Intercept) | 5.49 | 0.25 | 22.38 | 0.00 | |
| | | Cultivar (MM) | -0.15 | 0.35 | -0.43 | 0.68 | |
| | | Clinostat (SM) | 0.55 | 0.37 | 1.46 | 0.17 | |
| | | Cultivar (MM)*Clinostat (SM) | 0.05 | 0.51 | 0.10 | 0.92 | |
| | 5 | (Intercept) | 3.88 | 0.19 | 20.53 | 0.00 | |
| | | Cultivar (MM) | -0.58 | 0.27 | -2.16 | 0.05 | |
| | | Clinostat (SM) | -0.68 | 0.27 | -2.52 | 0.03 | |
| | | Cultivar (MM)*Clinostat (SM) | 0.52 | 0.38 | 1.38 | 0.19 | |
| **Trait** | **Trial** | | **Estimate** | **Std. Erro** | **t value** | **Pr(>\|t\|)** | **Transformation** |
| **Root Length** | 1 | (Intercept) | 10.95 | 1.16 | 9.43 | 0.00 | None |
| | | Cultivar (MM) | 3.42 | 1.64 | 2.08 | 0.07 | |
| | | Clinostat (SM) | -6.20 | 1.77 | -3.50 | 0.01 | |
| | | Cultivar (MM)*Clinostat (SM) | -2.02 | 2.68 | -0.75 | 0.47 | |
| | 2 | (Intercept) | 10.30 | 1.27 | 8.09 | 0.00 | |
| | | Cultivar (MM) | 1.75 | 1.80 | 0.97 | 0.35 | |
| | | Clinostat (SM) | -5.67 | 1.80 | -3.15 | 0.01 | |
| | | Cultivar (MM)*Clinostat (SM) | -2.45 | 2.55 | -0.96 | 0.35 | |
| | 3 | (Intercept) | 3.36 | 0.53 | 6.34 | 0.00 | |
| | | Cultivar (MM) | 0.96 | 0.84 | 1.15 | 0.28 | |
| | | Clinostat (SM) | 1.26 | 0.70 | 1.79 | 0.11 | |
| | | Cultivar (MM)*Clinostat (SM) | -2.34 | 1.06 | -2.21 | 0.05 | |
| | 4 | (Intercept) | 5.85 | 0.51 | 11.38 | 0.00 | |
| | | Cultivar (MM) | 0.89 | 0.73 | 1.22 | 0.25 | |
| | | Clinostat (SM) | 2.10 | 0.78 | 2.68 | 0.02 | |
| | | Cultivar (MM)*Clinostat (SM) | -2.09 | 1.07 | -1.96 | 0.08 | |
| | 5 | (Intercept) | 2.33 | 0.27 | 8.68 | 0.00 | |
| | | Cultivar (MM) | -0.23 | 0.38 | -0.60 | 0.57 | |
| | | Clinostat (SM) | -1.47 | 0.60 | -2.46 | 0.05 | |
| | | Cultivar (MM)*Clinostat (SM) | | | | | |
| **Trait** | **Trial** | | **Estimate** | **Std. Erro** | **t value** | **Pr(>\|t\|)** | **Transformation** |

| | | | | | | | |
|---|---|---|---|---|---|---|---|
| **Dry Mass (%)** | 1 | (Intercept) | 10.76 | 0.26 | 41.28 | 0.00 | None |
| | | Cultivar (MM) | 0.04 | 0.37 | 0.11 | 0.91 | |
| | | Clinostat (SM) | 1.85 | 0.40 | 4.63 | 0.00 | |
| | | Cultivar (MM)*Clinostat (SM) | -0.12 | 0.60 | -0.20 | 0.85 | |
| | 2 | (Intercept) | 10.95 | 0.38 | 28.82 | 0.00 | |
| | | Cultivar (MM) | 0.10 | 0.54 | 0.19 | 0.85 | |
| | | Clinostat (SM) | 1.58 | 0.54 | 2.93 | 0.01 | |
| | | Cultivar (MM)*Clinostat (SM) | 0.28 | 0.76 | 0.37 | 0.72 | |
| | 3 | (Intercept) | 14.64 | 0.48 | 30.30 | 0.00 | |
| | | Cultivar (MM) | -0.28 | 0.68 | -0.41 | 0.69 | |
| | | Clinostat (SM) | -1.21 | 0.68 | -1.77 | 0.10 | |
| | | Cultivar (MM)*Clinostat (SM) | 0.62 | 0.97 | 0.64 | 0.53 | |
| | 4 | (Intercept) | 12.60 | 0.22 | 57.60 | 0.00 | |
| | | Cultivar (MM) | 0.23 | 0.31 | 0.73 | 0.48 | |
| | | Clinostat (SM) | 0.29 | 0.33 | 0.87 | 0.41 | |
| | | Cultivar (MM)*Clinostat (SM) | 0.23 | 0.46 | 0.50 | 0.63 | |
| | 5 | (Intercept) | 12.48 | 0.73 | 17.00 | 0.00 | |
| | | Cultivar (MM) | 1.35 | 1.04 | 1.30 | 0.22 | |
| | | Clinostat (SM) | 0.89 | 1.04 | 0.85 | 0.41 | |
| | | Cultivar (MM)*Clinostat (SM) | 0.84 | 1.47 | 0.57 | 0.58 | |

**Table S8. Pairwise comparison summary table for each plant trait within each trial.** For each plant trait and each trial, the estimated marginal means, standard error, degrees of freedom, lower confidence level, upper confidence level, and Tukey HSD groupings (Group) are shown for each main effect (clinostat or cultivar).

| Trait | Replication | Cultivar | Clinostat | Estimated Marginal Means | Standard Error | Degrees of Freedom | Lower Confidence Level | Upper Confidence Level | Group | Effect |
|---|---|---|---|---|---|---|---|---|---|---|
| Apical Bud Height | 1 | MM | | 9.96 | 0.49 | 9 | 8.64 | 11.29 | a | Cultivar |
| | | H7996 | | 13.68 | 0.44 | 9 | 12.51 | 14.84 | b | |
| | | | C | 12.46 | 0.40 | 9 | 11.38 | 13.54 | a | Clinostat |
| | | | SM | 11.18 | 0.52 | 9 | 9.78 | 12.57 | a | |
| | 2 | MM | | 9.39 | 0.86 | 12 | 7.20 | 11.58 | a | Cultivar |
| | | H7996 | | 13.21 | 0.86 | 12 | 11.02 | 15.40 | b | |
| | | | C | 13.05 | 0.86 | 12 | 10.86 | 15.24 | b | Clinostat |
| | | | SM | 9.55 | 0.86 | 12 | 7.36 | 11.74 | a | |
| | 3 | MM | | 8.23 | 0.59 | 12 | 6.72 | 9.73 | a | Cultivar |
| | | H7996 | | 11.38 | 0.59 | 12 | 9.87 | 12.88 | b | |
| | | | C | 8.34 | 0.59 | 12 | 6.83 | 9.84 | a | Clinostat |
| | | | SM | 11.26 | 0.59 | 12 | 9.76 | 12.77 | b | |
| | 4 | MM | | 11.64 | 0.52 | 11 | 10.29 | 12.98 | a | Cultivar |
| | | H7996 | | 17.67 | 0.56 | 11 | 16.22 | 19.12 | b | |
| | | | C | 14.76 | 0.52 | 11 | 13.42 | 16.11 | a | Clinostat |
| | | | SM | 14.54 | 0.56 | 11 | 13.09 | 15.99 | a | |
| | 5 | MM | | 5.96 | 0.36 | 12 | 5.03 | 6.89 | a | Cultivar |
| | | H7996 | | 9.30 | 0.36 | 12 | 8.37 | 10.23 | b | |
| | | | C | 8.29 | 0.36 | 12 | 7.36 | 9.22 | b | Clinostat |
| | | | SM | 6.98 | 0.36 | 12 | 6.04 | 7.91 | a | |
| **Trait** | **Replication** | **Cultivar** | **Clinostat** | **Estimated Marginal Means** | **Standard Error** | **Degrees of Freedom** | **Lower Confidence Level** | **Upper Confidence Level** | **Group** | **Effect** |
| Shoot Fresh Mass | 1 | MM | | 10.22 | 1.14 | 9 | 7.17 | 13.27 | a | Cultivar |
| | | H7996 | | 9.33 | 1.00 | 9 | 6.64 | 12.02 | a | |
| | | | C | 11.91 | 0.93 | 9 | 9.42 | 14.40 | b | Clinostat |
| | | | SM | 7.63 | 1.20 | 9 | 4.42 | 10.85 | a | |
| | 2 | MM | | 9.64 | 0.60 | 12 | 8.10 | 11.18 | a | Cultivar |
| | | H7996 | | 8.53 | 0.60 | 12 | 6.99 | 10.07 | a | |
| | | | C | 11.58 | 0.60 | 12 | 10.04 | 13.13 | b | Clinostat |
| | | | SM | 6.59 | 0.60 | 12 | 5.04 | 8.13 | a | |
| | 3 | MM | | 4.97 | 0.45 | 12 | 3.82 | 6.11 | a | Cultivar |
| | | H7996 | | 5.17 | 0.45 | 12 | 4.02 | 6.32 | a | |
| | | | C | 3.08 | 0.45 | 12 | 1.93 | 4.23 | a | Clinostat |
| | | | SM | 7.06 | 0.45 | 12 | 5.91 | 8.21 | b | |
| | 4 | MM | | 10.93 | 0.61 | 11 | 9.35 | 12.51 | a | Cultivar |
| | | H7996 | | 10.69 | 0.66 | 11 | 8.99 | 12.40 | a | |
| | | | C | 10.21 | 0.61 | 11 | 8.63 | 11.79 | a | Clinostat |
| | | | SM | 11.41 | 0.66 | 11 | 9.70 | 13.12 | a | |
| | 5 | MM | | 2.04 | 0.23 | 12 | 1.45 | 2.63 | a | Cultivar |
| | | H7996 | | 2.61 | 0.23 | 12 | 2.02 | 3.20 | a | |
| | | | C | 2.94 | 0.23 | 12 | 2.35 | 3.53 | b | Clinostat |
| | | | SM | 1.71 | 0.23 | 12 | 1.12 | 2.30 | a | |
| **Trait** | **Replication** | **Cultivar** | **Clinostat** | **Estimated Marginal Means** | **Standard Error** | **Degrees of Freedom** | **Lower Confidence Level** | **Upper Confidence Level** | **Group** | **Effect** |
| | 1 | MM | | 1.17 | 0.13 | 9 | 0.83 | 1.51 | a | Cultivar |
| | | H7996 | | 1.07 | 0.11 | 9 | 0.78 | 1.37 | a | |

| | | | | | | | | | | |
|---|---|---|---|---|---|---|---|---|---|---|
| | 1 | | C | 1.28 | 0.10 | 9 | 1.01 | 1.56 | a | Clinostat |
| | | | SM | 0.96 | 0.13 | 9 | 0.60 | 1.32 | a | |
| | 2 | MM | | 1.13 | 0.07 | 12 | 0.96 | 1.30 | a | Cultivar |
| | | H7996 | | 0.98 | 0.07 | 12 | 0.81 | 1.15 | a | |
| | | | C | 1.27 | 0.07 | 12 | 1.10 | 1.44 | b | Clinostat |
| | | | SM | 0.84 | 0.07 | 12 | 0.67 | 1.01 | a | |
| **Shoot Dry Mass** | 3 | MM | | 0.69 | 0.07 | 12 | 0.51 | 0.88 | a | Cultivar |
| | | H7996 | | 0.72 | 0.07 | 12 | 0.53 | 0.91 | a | |
| | | | C | 0.44 | 0.07 | 12 | 0.26 | 0.63 | a | Clinostat |
| | | | SM | 0.97 | 0.07 | 12 | 0.78 | 1.15 | b | |
| | 4 | MM | | 1.43 | 0.08 | 11 | 1.22 | 1.64 | a | Cultivar |
| | | H7996 | | 1.36 | 0.09 | 11 | 1.14 | 1.59 | a | |
| | | | C | 1.30 | 0.08 | 11 | 1.09 | 1.51 | a | Clinostat |
| | | | SM | 1.50 | 0.09 | 11 | 1.27 | 1.72 | a | |
| | 5 | MM | | 0.29 | 0.03 | 12 | 0.22 | 0.36 | a | Cultivar |
| | | H7996 | | 0.33 | 0.03 | 12 | 0.26 | 0.40 | a | |
| | | | C | 0.38 | 0.03 | 12 | 0.31 | 0.45 | b | Clinostat |
| | | | SM | 0.25 | 0.03 | 12 | 0.17 | 0.32 | a | |

| Trait | Replication | Cultivar | Clinostat | Estimated Marginal Means | Standard Error | Degrees of Freedom | Lower Confidence Level | Upper Confidence Level | Group | Effect |
|---|---|---|---|---|---|---|---|---|---|---|
| | 1 | MM | | 5.40 | 0.19 | 9 | 4.88 | 5.91 | a | Cultivar |
| | | H7996 | | 5.35 | 0.17 | 9 | 4.89 | 5.80 | a | |
| | | | C | 5.64 | 0.16 | 9 | 5.22 | 6.06 | a | Clinostat |
| | | | SM | 5.10 | 0.20 | 9 | 4.56 | 5.65 | a | |
| | 2 | MM | | 5.99 | 0.16 | 12 | 5.58 | 6.40 | b | Cultivar |
| | | H7996 | | 5.37 | 0.16 | 12 | 4.96 | 5.78 | a | |
| | | | C | 5.97 | 0.16 | 12 | 5.56 | 6.38 | b | Clinostat |
| | | | SM | 5.39 | 0.16 | 12 | 4.98 | 5.80 | a | |
| **Shoot Diameter** | 3 | MM | | 4.35 | 0.11 | 12 | 4.07 | 4.63 | a | Cultivar |
| | | H7996 | | 4.46 | 0.11 | 12 | 4.18 | 4.74 | a | |
| | | | C | 3.66 | 0.11 | 12 | 3.38 | 3.93 | a | Clinostat |
| | | | SM | 5.15 | 0.11 | 12 | 4.88 | 5.43 | b | |
| | 4 | MM | | 5.64 | 0.17 | 11 | 5.19 | 6.09 | a | Cultivar |
| | | H7996 | | 5.76 | 0.19 | 11 | 5.28 | 6.25 | a | |
| | | | C | 5.42 | 0.17 | 11 | 4.97 | 5.87 | a | Clinostat |
| | | | SM | 5.99 | 0.19 | 11 | 5.50 | 6.47 | b | |
| | 5 | MM | | 3.23 | 0.13 | 12 | 2.89 | 3.57 | a | Cultivar |
| | | H7996 | | 3.55 | 0.13 | 12 | 3.20 | 3.89 | a | |
| | | | SM | 3.18 | 0.13 | 12 | 2.84 | 3.52 | a | Clinostat |
| | | | C | 3.59 | 0.13 | 12 | 3.25 | 3.93 | b | |

| Trait | Replication | Cultivar | Clinostat | Estimated Marginal Means | Standard Error | Degrees of Freedom | Lower Confidence Level | Upper Confidence Level | Group | Effect |
|---|---|---|---|---|---|---|---|---|---|---|
| | 1 | MM | | 26.05 | 2.55 | 9 | 19.21 | 32.89 | a | Cultivar |
| | | H7996 | | 19.93 | 2.25 | 9 | 13.91 | 25.96 | a | |
| | | | C | 32.15 | 2.08 | 9 | 26.57 | 37.73 | b | Clinostat |
| | | | SM | 13.83 | 2.69 | 9 | 6.63 | 21.04 | a | |
| | 2 | MM | | 20.30 | 2.29 | 12 | 14.46 | 26.15 | a | Cultivar |
| | | H7996 | | 18.97 | 2.29 | 12 | 13.12 | 24.81 | a | |
| | | | C | 28.40 | 2.29 | 12 | 22.55 | 34.24 | b | Clinostat |
| | | | SM | 10.87 | 2.29 | 12 | 5.03 | 16.72 | a | |

| | | | | | | | | | | |
|---|---|---|---|---|---|---|---|---|---|---|
| Root Length | 3 | MM | | 6.86 | 1.53 | 12 | 2.95 | 10.76 | a | Cultivar |
| | | H7996 | | 9.06 | 1.53 | 12 | 5.16 | 12.97 | a | |
| | | | C | 5.94 | 1.53 | 12 | 2.04 | 9.85 | a | Clinostat |
| | | | SM | 9.98 | 1.53 | 12 | 6.07 | 13.88 | a | |
| | 4 | MM | | 17.11 | 0.92 | 11 | 14.73 | 19.50 | a | Cultivar |
| | | H7996 | | 17.52 | 1.00 | 11 | 14.94 | 20.10 | a | |
| | | | C | 15.98 | 0.92 | 11 | 13.59 | 18.36 | a | Clinostat |
| | | | SM | 18.66 | 1.00 | 11 | 16.08 | 21.23 | a | |
| | 5 | MM | | 2.67 | 0.39 | 12 | 1.67 | 3.67 | a | Cultivar |
| | | H7996 | | 3.23 | 0.39 | 12 | 2.23 | 4.23 | a | |
| | | | C | 5.63 | 0.39 | 12 | 4.63 | 6.62 | b | Clinostat |
| | | | SM | 0.27 | 0.39 | 12 | -0.73 | 1.27 | a | |

| Trait | Replication | Cultivar | Clinostat | Estimated Marginal Means | Standard Error | Degrees of Freedom | Lower Confidence Level | Upper Confidence Level | Group | Effect |
|---|---|---|---|---|---|---|---|---|---|---|
| Dry Mass (%) | 1 | MM | | 11.66 | 0.23 | 9 | 11.06 | 12.27 | a | Cultivar |
| | | H7996 | | 11.68 | 0.20 | 9 | 11.15 | 12.21 | a | |
| | | | C | 10.78 | 0.18 | 9 | 10.29 | 11.27 | a | Clinostat |
| | | | SM | 12.56 | 0.24 | 9 | 11.93 | 13.20 | b | |
| | 2 | MM | | 11.99 | 0.27 | 12 | 11.30 | 12.67 | a | Cultivar |
| | | H7996 | | 11.74 | 0.27 | 12 | 11.06 | 12.43 | a | |
| | | | C | 11.01 | 0.27 | 12 | 10.32 | 11.69 | a | Clinostat |
| | | | SM | 12.72 | 0.27 | 12 | 12.04 | 13.41 | b | |
| | 3 | MM | | 14.07 | 0.34 | 12 | 13.19 | 14.94 | a | Cultivar |
| | | H7996 | | 14.04 | 0.34 | 12 | 13.16 | 14.91 | a | |
| | | | C | 14.50 | 0.34 | 12 | 13.63 | 15.37 | a | Clinostat |
| | | | SM | 13.60 | 0.34 | 12 | 12.73 | 14.47 | a | |
| | 4 | MM | | 13.09 | 0.15 | 11 | 12.69 | 13.49 | a | Cultivar |
| | | H7996 | | 12.75 | 0.17 | 11 | 12.32 | 13.18 | a | |
| | | | C | 12.72 | 0.15 | 11 | 12.32 | 13.12 | a | Clinostat |
| | | | SM | 13.12 | 0.17 | 11 | 12.69 | 13.55 | a | |
| | 5 | MM | | 14.70 | 0.52 | 12 | 13.38 | 16.03 | b | Cultivar |
| | | H7996 | | 12.93 | 0.52 | 12 | 11.60 | 14.25 | a | |
| | | | C | 13.16 | 0.52 | 12 | 11.83 | 14.49 | a | Clinostat |
| | | | SM | 14.47 | 0.52 | 12 | 13.14 | 15.80 | a | |

**Table S9.** ANOVA table for each plant trait within upright control conditions. To determine if significant differences were observed between cultivar or trial, a two-way analysis of variance (ANOVA) was used for each trait within upright control conditions. Within each model, cultivar and trial were included as independent variables.

| | | Df | Sum Sq | Mean Sq | F value | Pr(>F) |
|---|---|---|---|---|---|---|
| | Cultivar | 1 | 152.88 | 152.88 | 61.25 | 4.14E-09 |
| **ical Bud Hei** | Trial | 4 | 273.78 | 68.44 | 27.42 | 3.19E-10 |
| | Residuals | 34 | 84.86 | 2.50 | NA | NA |

| | | Df | Sum Sq | Mean Sq | F value | Pr(>F) |
|---|---|---|---|---|---|---|
| | Cultivar | 1 | 6.34 | 6.34 | 1.38 | 2.49E-01 |
| **oot Fresh M** | Trial | 4 | 662.70 | 165.68 | 35.95 | 9.03E-12 |
| | Residuals | 34 | 156.71 | 4.61 | NA | NA |

| | | Df | Sum Sq | Mean Sq | F value | Pr(>F) |
|---|---|---|---|---|---|---|
| | Cultivar | 1 | 0.09 | 0.09 | 1.52 | 2.26E-01 |
| **hoot Dry Ma** | Trial | 4 | 7.36 | 1.84 | 30.04 | 9.86E-11 |
| | Residuals | 34 | 2.08 | 0.06 | NA | NA |

| | | Df | Sum Sq | Mean Sq | F value | Pr(>F) |
|---|---|---|---|---|---|---|
| | Cultivar | 1 | 0.00 | 0.00 | 0.02 | 8.91E-01 |
| **hoot Diamet** | Trial | 4 | 41.62 | 10.40 | 40.01 | 2.08E-12 |
| | Residuals | 34 | 8.84 | 0.26 | NA | NA |

| | | Df | Sum Sq | Mean Sq | F value | Pr(>F) |
|---|---|---|---|---|---|---|
| | Cultivar | 1 | 77.28 | 77.28 | 2.41 | 1.30E-01 |
| **Root Length** | Trial | 4 | 4880.94 | 1220.23 | 38.01 | 4.21E-12 |
| | Residuals | 34 | 1091.46 | 32.10 | NA | NA |

**Table S10. Pairwise comparison summary table for each plant trait in upright control conditions.** For each plant trait, summary statistics and Tukey HSD groupings (Group) are shown for each trial in upright control conditions.

| | Trial | Mean | Standard D | N | Standard E | Min | Max | Q25 | Q50 | Q75 | Group |
|---|---|---|---|---|---|---|---|---|---|---|---|
| **Apical Bud Height** | February | 12.46 | 2.01 | 8 | 0.56 | 9.30 | 15.30 | 11.03 | 13.15 | 13.58 | b |
| | March | 13.05 | 2.76 | 8 | 0.56 | 9.20 | 17.40 | 11.33 | 12.80 | 14.88 | ab |
| | May | 14.76 | 3.05 | 8 | 0.56 | 9.80 | 18.70 | 13.38 | 15.35 | 16.75 | a |
| | April | 8.34 | 2.75 | 8 | 0.56 | 5.40 | 13.10 | 6.38 | 7.65 | 9.20 | c |
| | June | 8.29 | 2.33 | 8 | 0.56 | 4.90 | 11.20 | 6.80 | 8.55 | 9.68 | c |

| | Trial | Mean | Standard D | N | Standard E | Min | Max | Q25 | Q50 | Q75 | Group |
|---|---|---|---|---|---|---|---|---|---|---|---|
| **Shoot Fresh Mass** | February | 11.91 | 3.25 | 8 | 0.76 | 8.59 | 17.51 | 9.47 | 10.87 | 14.38 | a |
| | March | 11.58 | 2.56 | 8 | 0.76 | 8.93 | 14.97 | 9.56 | 10.77 | 14.11 | a |
| | May | 10.21 | 2.00 | 8 | 0.76 | 7.75 | 13.08 | 8.84 | 9.51 | 11.83 | a |
| | April | 3.08 | 1.15 | 8 | 0.76 | 1.75 | 4.65 | 2.26 | 2.59 | 4.11 | b |
| | June | 2.94 | 0.95 | 8 | 0.76 | 1.48 | 3.86 | 2.37 | 3.26 | 3.64 | b |

| | Trial | Mean | Standard D | N | Standard E | Min | Max | Q25 | Q50 | Q75 | Group |
|---|---|---|---|---|---|---|---|---|---|---|---|
| **Shoot Dry Mass** | February | 1.28 | 0.36 | 8 | 0.09 | 0.92 | 1.96 | 1.00 | 1.20 | 1.50 | a |
| | March | 1.27 | 0.28 | 8 | 0.09 | 0.96 | 1.76 | 1.06 | 1.20 | 1.45 | a |
| | May | 1.30 | 0.26 | 8 | 0.09 | 0.99 | 1.72 | 1.10 | 1.26 | 1.46 | a |
| | April | 0.44 | 0.16 | 8 | 0.09 | 0.27 | 0.67 | 0.34 | 0.37 | 0.56 | b |
| | June | 0.38 | 0.10 | 8 | 0.09 | 0.20 | 0.50 | 0.32 | 0.41 | 0.45 | b |

| | Trial | Mean | Standard D | N | Standard E | Min | Max | Q25 | Q50 | Q75 | Group |
|---|---|---|---|---|---|---|---|---|---|---|---|
| **Shoot Diameter** | February | 5.64 | 0.57 | 8 | 0.18 | 4.76 | 6.30 | 5.22 | 5.80 | 6.07 | a |
| | March | 5.97 | 0.47 | 8 | 0.18 | 5.35 | 6.47 | 5.56 | 6.06 | 6.41 | a |
| | May | 5.42 | 0.51 | 8 | 0.18 | 4.68 | 6.14 | 5.14 | 5.49 | 5.74 | a |
| | April | 3.66 | 0.45 | 8 | 0.18 | 3.01 | 4.32 | 3.30 | 3.68 | 4.00 | b |
| | June | 3.59 | 0.51 | 8 | 0.18 | 2.74 | 4.38 | 3.39 | 3.63 | 3.84 | b |

| | Trial | Mean | Standard D | N | Standard E | Min | Max | Q25 | Q50 | Q75 | Group |
|---|---|---|---|---|---|---|---|---|---|---|---|
| **Root Length** | February | 32.15 | 8.04 | 8 | 2.00 | 21.55 | 46.70 | 28.18 | 33.33 | 35.01 | a |
| | March | 28.40 | 7.98 | 8 | 2.00 | 18.30 | 41.73 | 23.36 | 27.01 | 33.58 | a |
| | May | 15.98 | 2.50 | 8 | 2.00 | 13.77 | 20.47 | 14.33 | 14.84 | 17.06 | b |
| | April | 5.94 | 5.55 | 8 | 2.00 | 0.00 | 13.66 | 0.00 | 6.43 | 10.20 | c |
| | June | 5.63 | 1.30 | 8 | 2.00 | 3.13 | 7.42 | 5.04 | 5.71 | 6.43 | c |

**Table S11.** ANOVA table for each plant trait within simulated microgravity conditions. To determine if significant differences were observed between cultivar or trial, a two-way analysis of variance (ANOVA) was used for each trait within simulated microgravity conditions. Within each model, cultivar and trial were included as independent variables. A Tukey's ladder of powers transformation was used if residuals were not from a normal distribution.

| | | Df | Sum Sq | Mean Sq | F value | Pr(>F) | Transformation |
|---|---|---|---|---|---|---|---|
| | Cultivar | 1 | 0.23 | 0.23 | 39.83 | 5.87E-07 | |
| **Apical Bud Height** | Replication | 4 | 0.42 | 0.10 | 18.18 | 1.12E-07 | Tukey |
| | Residuals | 30 | 0.17 | 0.01 | | | |

| | | Df | Sum Sq | Mean Sq | F value | Pr(>F) | Transformation |
|---|---|---|---|---|---|---|---|
| | Cultivar | 1 | 0.02 | 0.02 | 0.03 | 8.62E-01 | |
| **Shoot Fresh Mass** | Replication | 4 | 360.08 | 90.02 | 123.70 | 3.44E-18 | None |
| | Residuals | 30 | 21.83 | 0.73 | | | |

| | | Df | Sum Sq | Mean Sq | F value | Pr(>F) | Transformation |
|---|---|---|---|---|---|---|---|
| | Cultivar | 1 | 0.01 | 0.01 | 0.57 | 4.57E-01 | |
| **Shoot Dry Mass** | Replication | 4 | 6.01 | 1.50 | 74.91 | 3.56E-15 | None |
| | Residuals | 30 | 0.60 | 0.02 | | | |

| | | Df | Sum Sq | Mean Sq | F value | Pr(>F) | Transformation |
|---|---|---|---|---|---|---|---|
| | Cultivar | 1 | 0.20 | 0.20 | 0.89 | 3.53E-01 | |
| **Shoot Diameter** | Replication | 4 | 34.55 | 8.64 | 37.88 | 2.53E-11 | None |
| | Residuals | 30 | 6.84 | 0.23 | | | |

| | | Df | Sum Sq | Mean Sq | F value | Pr(>F) | Transformation |
|---|---|---|---|---|---|---|---|
| | Cultivar | 1 | 11.51 | 11.51 | 1.66 | 2.08E-01 | |
| **Root Length** | Replication | 4 | 1327.84 | 331.96 | 47.75 | 1.37E-12 | None |
| | Residuals | 30 | 208.55 | 6.95 | | | |

**Table S12. Pairwise comparison summary table for each plant trait in simulated microgravity conditions.** For each plant trait, summary statistics and Tukey HSD groupings (Group) are shown for each trial in simulated microgravity conditions.

| | Trial | Mean | Standard D | N | Standard E | Min | Max | Q25 | Q50 | Q75 | Group |
|---|---|---|---|---|---|---|---|---|---|---|---|
| | February | 1.84 | 0.10 | 5 | 0.03 | 1.70 | 1.93 | 1.76 | 1.91 | 1.91 | ab |
| | March | 1.74 | 0.15 | 8 | 0.03 | 1.54 | 1.95 | 1.65 | 1.69 | 1.87 | b |
| **Apical Bud Height** | May | 1.92 | 0.14 | 7 | 0.03 | 1.76 | 2.09 | 1.83 | 1.86 | 2.05 | a |
| | April | 1.83 | 0.07 | 8 | 0.03 | 1.70 | 1.92 | 1.81 | 1.83 | 1.87 | ab |
| | June | 1.62 | 0.10 | 8 | 0.03 | 1.49 | 1.76 | 1.54 | 1.62 | 1.69 | c |

| | Trial | Mean | Standard D | N | Standard E | Min | Max | Q25 | Q50 | Q75 | Group |
|---|---|---|---|---|---|---|---|---|---|---|---|
| | February | 7.72 | 0.88 | 5 | 0.38 | 6.21 | 8.40 | 7.76 | 8.00 | 8.22 | b |
| | March | 6.59 | 0.28 | 8 | 0.30 | 6.09 | 6.96 | 6.44 | 6.60 | 6.80 | b |
| **Shoot Fresh Mass** | May | 11.39 | 0.99 | 7 | 0.32 | 10.55 | 13.10 | 10.60 | 10.89 | 12.01 | a |
| | April | 7.06 | 1.31 | 8 | 0.30 | 5.25 | 8.55 | 6.34 | 6.88 | 8.40 | b |
| | June | 1.71 | 0.25 | 8 | 0.30 | 1.23 | 2.01 | 1.61 | 1.70 | 1.88 | c |

| | Trial | Mean | Standard D | N | Standard E | Min | Max | Q25 | Q50 | Q75 | Group |
|---|---|---|---|---|---|---|---|---|---|---|---|
| | February | 0.97 | 0.13 | 5 | 0.06 | 0.77 | 1.13 | 0.93 | 0.99 | 1.04 | b |
| | March | 0.84 | 0.06 | 8 | 0.05 | 0.75 | 0.91 | 0.80 | 0.84 | 0.89 | b |
| **Shoot Dry Mass** | May | 1.50 | 0.14 | 7 | 0.05 | 1.35 | 1.76 | 1.42 | 1.43 | 1.56 | a |
| | April | 0.97 | 0.23 | 8 | 0.05 | 0.62 | 1.26 | 0.87 | 0.93 | 1.18 | b |
| | June | 0.25 | 0.04 | 8 | 0.05 | 0.19 | 0.31 | 0.23 | 0.25 | 0.26 | c |

| | Trial | Mean | Standard D | N | Standard E | Min | Max | Q25 | Q50 | Q75 | Group |
|---|---|---|---|---|---|---|---|---|---|---|---|
| | February | 5.19 | 0.73 | 5 | 0.21 | 4.43 | 6.41 | 4.95 | 5.06 | 5.10 | ab |
| | March | 5.39 | 0.59 | 8 | 0.17 | 4.45 | 6.31 | 5.12 | 5.37 | 5.56 | ab |
| **Shoot Diameter** | May | 5.98 | 0.38 | 7 | 0.18 | 5.44 | 6.39 | 5.66 | 6.20 | 6.26 | a |
| | April | 5.15 | 0.36 | 8 | 0.17 | 4.56 | 5.71 | 4.95 | 5.22 | 5.34 | b |
| | June | 3.18 | 0.28 | 8 | 0.17 | 2.80 | 3.60 | 3.00 | 3.20 | 3.36 | c |

| | Trial | Mean | Standard D | N | Standard E | Min | Max | Q25 | Q50 | Q75 | Group |
|---|---|---|---|---|---|---|---|---|---|---|---|
| | February | 13.48 | 2.58 | 5 | 1.18 | 9.80 | 16.11 | 11.88 | 14.50 | 15.11 | b |
| | March | 10.87 | 3.84 | 8 | 0.93 | 8.51 | 19.52 | 8.76 | 9.01 | 10.90 | b |
| **Root Length** | May | 18.44 | 3.09 | 7 | 1.00 | 14.64 | 21.69 | 16.11 | 17.78 | 21.36 | a |
| | April | 9.98 | 2.23 | 8 | 0.93 | 7.62 | 13.41 | 8.39 | 9.18 | 11.95 | b |
| | June | 0.27 | 0.77 | 8 | 0.93 | 0.00 | 2.17 | 0.00 | 0.00 | 0.00 | c |
